# Dopaminergic axons track somatic signaling in behaving mice

**DOI:** 10.1101/2022.06.20.496872

**Authors:** Maite Azcorra, Zachary Gaertner, Connor Davidson, Charu Ramakrishnan, Lief Fenno, Yoon Seok Kim, Karl Deisseroth, Rajeshwar Awatramani, Daniel A. Dombeck

## Abstract

Striatal dopamine released from the axons of midbrain dopamine neurons has been linked to a wide range of functions, including movement control and reward-based learning. Recent studies have reported functional signaling differences between axons and somas of dopamine neurons, suggesting that local modulation controls dopamine release and calling into question the classical view of somatic control. However, these experiments are technically challenging, making it difficult to ensure that axonal and somatic recordings come from the same neurons, particularly given the heterogeneity of dopaminergic cell types. Here we used genetic strategies to isolate key dopaminergic neuron subtypes and monitor their axonal and somatic signaling patterns in behaving mice. Contrary to the inferences drawn from previous studies, these experiments revealed a robust correlation between somatic and axonal signaling. Thus, by exploiting a previously unknown connection between genetic and functional diversity in dopamine neurons, we establish that subtypes must be considered to understand the mechanisms of dopamine release in striatum during behavior.

---

Midbrain dopamine neurons play a role in a wide range of behaviors^1–9^. For example, dopamine neurons of the VTA and SNc fire bursts of action potentials in response to unexpected rewards^1^, and SNc neuron firing correlates with movement and motor learning^2–7^. Furthermore, optogenetic perturbations of dopamine neurons can causally impact reward-based learning^10,11^ and motor behavior^3,4,12^. A major target of these midbrain neurons is the striatum, where dopamine is released by their axons. However, it is currently unclear whether this dopamine release is driven mainly by somatic firing or by local striatal mechanisms. For decades, it was assumed that striatal dopamine release was controlled by anterogradely propagating action potentials originating in the midbrain somas; but this classical view has recently been called into question by research demonstrating the ability of local mechanisms to control striatal dopamine release independently of midbrain somatic firing^13–17^.

Mechanistically, *in vitro* studies have shown that coordinated activation of striatal cholinergic interneurons can not only modulate, but also trigger dopamine release in the absence of somatic firing^13–16^. Pioneering *in vivo* studies have recently provided strong support for the idea that this local mechanism plays a significant role in dopamine release during behavior. One study recorded striatal dopamine and acetylcholine release dynamics (using fluorescent sensors and photometry recordings) during movement in freely behaving mice and found that dopamine and acetylcholine signaling co-varied with mouse movement direction, and that inhibition of nicotinic receptors in striatum decreased dopamine signaling^16^. Another study recorded VTA cell body firing (with single-unit recording) and striatal dopamine release (with photometry of a fluorescent dopamine sensor and microdialysis) during a reward-based choice task and found that dopamine release from striatal axons co-varied with reward expectation, while firing in the midbrain somas did not^17^. This study further observed fast striatal dopamine release during certain behavioral epochs that did not correspond with somatic firing, suggesting these differences were due to local modulation. Thus, cholinergic mechanisms can drive dopamine release independently of somatic firing, and both *in vivo* covariations of dopamine-acetylcholine signaling and differences between midbrain somatic firing and striatal dopamine release dynamics support the idea that these mechanisms are relevant during behavior.

However, establishing that dopamine is released from axons independently of somatic firing *in vivo* requires that axonal and somatic recordings are made from the same neurons. Thus, an alternative explanation for the observed soma-axon signaling differences is that the striatal dopamine detected was released by a different set of axons than those belonging to the recorded somas. This point may be especially important if different functional subtypes of dopaminergic neurons exist with intermingled somas in the midbrain and axons in the striatum. However, evidence for the existence of functional subtypes of dopamine neurons has been controversial^18^ and complicated by the lack of labeling strategies to isolate and record from such subtypes.

Here we used genetic strategies to isolate dopaminergic subtypes for labeling and recording. We used fiber photometry to record GCaMP calcium transients from both axons and somas of isolated dopaminergic subtypes in head-fixed mice running on a treadmill. GCaMP is ideally suited for these experiments because all known mechanisms for triggering axonal dopamine release involve increases in intracellular calcium concentration^19,20^, including anterogradely propagating action potentials^21–25^ and cholinergic modulation^14,16^. At the soma, GCaMP transients are caused by somatic action potential firing, thus enabling the use of a single strategy to measure a proxy of dopamine release from axons and firing from somas. Critically, the detected calcium transients are generated only from the labeled neurons; non-labeled neurons do not contribute. For this reason, GCaMP is preferred over extracellular dopamine sensors (i.e. dLight, GRAB-DA, microdialysis) because axons from different genetic subtypes densely overlap in many striatal regions^26^ and these sensors detect dopamine released from all nearby axons, without subtype specificity.

Locomotion is also well suited for investigating potential soma-axon signaling differences since it is known to be associated with rapid signaling in dopaminergic axons^3^, somas^4,7^, and cholinergic interneurons^16,27,28^. Though treadmill locomotion is similar to the freely moving behavior used in one previous *in vivo* study that found evidence for local control of striatal dopamine release^16^, it is quite different from the reward/choice-based paradigm from the other study^17^. Because of these behavioral and recording method differences, before recording from dopamine neuron subtypes, we first asked whether we could reproduce the soma-axon signaling differences previously described in non-subtype specific recordings, but with GCaMP and head-fixed locomotion.

We labelled non-subtype-specific DAT-expressing neurons in SNc with GCaMP6f (DAT-Cre mouse + AAV-FLEX-GCaMP6f), and used fiber photometry to simultaneously record from populations of axons in the striatum with one fiber and SNc somas with another fiber (Fig. 1b-c) while head-fixed mice ran on a treadmill. To control for any movement artifacts, we also recorded GCaMP6f fluorescence at its isosbestic wavelength, 405 nm^29^. We recorded from a broad range of locations within striatum and SNc, and observed highly dissimilar signaling (ΔF/F) between striatal axons and SNc somas (Fig. 1d) across recordings and mice. On average, the cross-correlation between soma and axon ΔF/F was relatively low (mean = 0.37), indicative of weakly correlated signaling patterns (Fig. 1e-g). Therefore, similarly to previous reports^17^, we find somatic and axonal dopamine neuron signaling that are often very different when dopamine neurons are indiscriminately labelled. This is consistent with the hypothesis that local cholinergic modulation drives axonal signaling independent of somatic firing during locomotion^16^.

**Figure 1:**
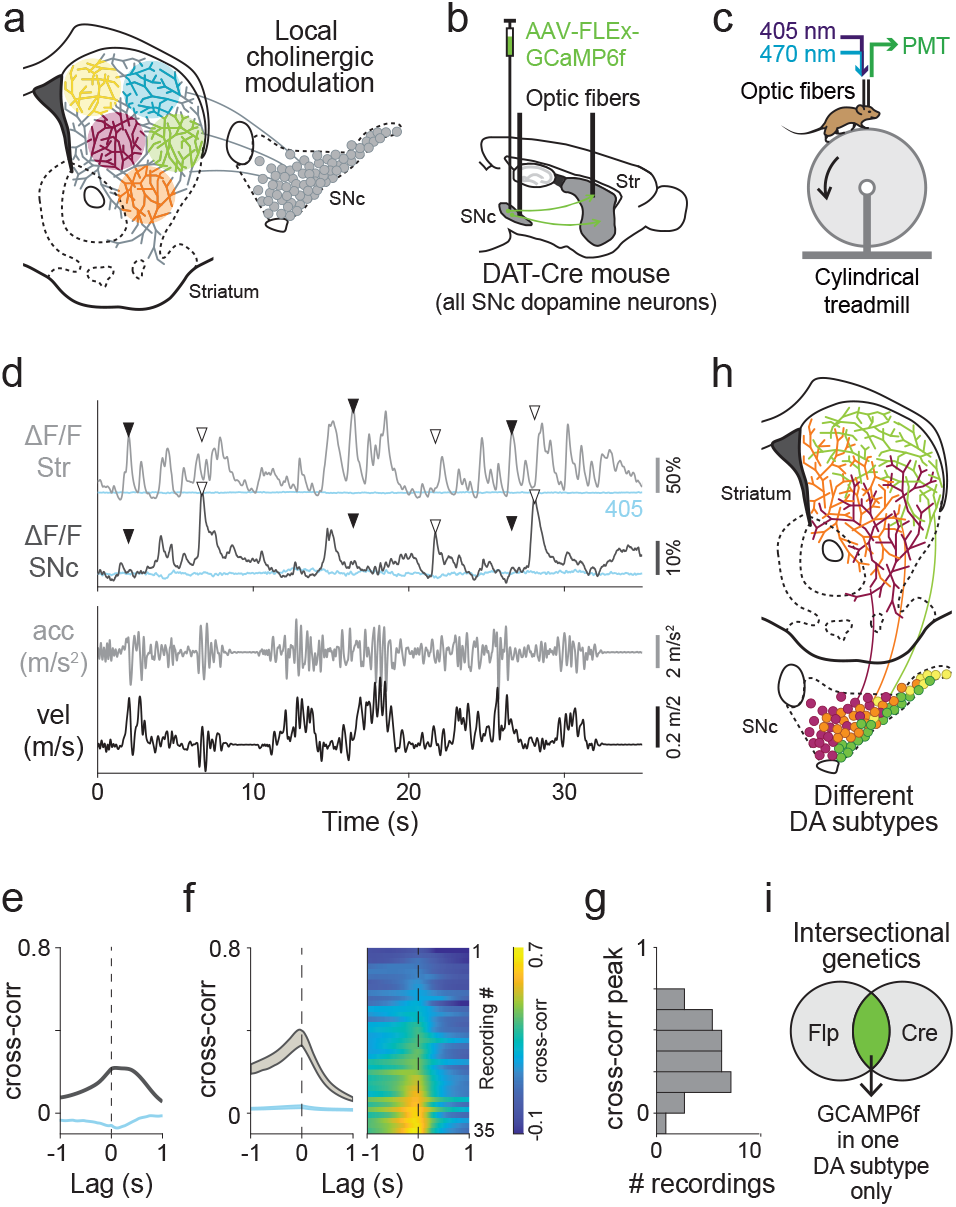
Differences in calcium transients between somas and axons of striatonigral dopaminergic neurons could result from local neuromodulation and/or overlapping dopamine neuron subtypes with different functions. a. Cholinergic interneurons could drive dopamine release from dopamine axons locally in striatum independently of dopamine soma firing, resulting in discrepancies between soma and axon signaling during behavior. b. Schematic showing experimental design and recording locations for d-g. c. Head-fixed mouse running on treadmill during fiber photometry, with simultaneous recording of calcium transients (470 nm excitation) and isosbestic control (405 nm excitation). d. Example velocity, acceleration, and calcium ΔF/F traces from simultaneous recording from dopamine axons in striatum and somas in SNc in a mouse where SNc dopamine neurons are indiscriminately labeled. Isosbestic control shown in blue. ▾= example transients present in striatum but not in SNc, ▽ = example transients present in SNc but not in striatum. e. Cross-correlation between simultaneous ΔF/F traces in striatum and SNc for traces shown in d. Isosbestic control shown in blue. f. Average cross-correlation between simultaneous ΔF/F traces in striatum and SNc from multiple recordings and mice. Isosbestic control shown in blue. Shaded regions denote mean ± s.e.m. Heatmap shows cross correlations for each recording sorted by peak magnitude (mice = 5, n = 35 recordings). g. Histogram of peak cross-correlations between SNc and Str for all recordings shown in f. h. Different functional subtypes of dopamine neurons could also explain differences in signaling between axons and somas, depending on the recording locations. i. Schematic of intersectional genetic strategy used to isolate single dopamine subtypes and label them with GCaMP6f, as used in figures 2-4.

However, these results could also be explained by different functional dopaminergic subtypes with intermingled somas in the midbrain and axons in the striatum (Fig. 1h)—but testing this hypothesis requires the ability to uniquely and consistently label and record from these functional subtypes, which has proven difficult. We and others recently described several dopamine neuron subtypes defined by their molecular expression profiles^30,31^, which we showed have diverse, but somewhat overlapping, cell body locations in SNc and projection targets in striatum^26^. We reasoned that these genetic subtypes may also have different functional signaling patterns and, if so, we could test whether isolating them leads to more similar soma-axon signaling. We focused on three SNc genetic subtypes that account for most of the dopamine neurons in SNc: Aldh1a1+ (somas in ventral SNc, axons in dorso-lateral striatum), Calb1+ (dorsal SNc, ventral and medial striatum), and VGlut2+ (lateral SNc, posterior striatum) ^26^ (Fig. S1b). To isolate each subtype, we used our previously described intersectional genetic strategies, labelling only neurons expressing a dopamine specific gene and a subtype-specific marker gene^26,32,33^ (Fig. 1i, Fig. S1a). For isolating Aldh1a1+ dopamine neurons, we generated a new Cre line that allows for more comprehensive and robust labelling compared to our previous tamoxifen-dependent line (Fig. S2).

We first used fiber photometry to determine whether striatal axons from different genetic subtypes differ in their functional signaling patterns during locomotion (Fig. 2a-c). In individual recordings, we observed that Calb1+ and VGlut2+ axons preferentially signaled during locomotion decelerations, while Aldh1a1+ axons preferentially signaled during locomotion accelerations (Fig. 2c’). Accordingly, cross-correlations between calcium ΔF/F traces (ΔF/F traces) and acceleration revealed a deep trough at positive time lags for Calb1+ and VGlut2+ axons (which is indicative of calcium transients following decelerations), but a large peak at positive lags for Aldh1a1+ axons (transients following accelerations) (Fig. 2c’’-f), when recorded from a wide range of locations within striatum (Fig. 2e,f). To further quantify these differences, we used a dimensionality reduction technique to extract the components that best explain the variance in the cross-correlations. We applied principal component analysis (PCA) to the matrix of all cross-correlation traces from all subtypes (see Methods), finding that the first principal component (PC1) explained a large amount (63%) of the variance. Positive PC1 (+PC1) closely approximated the average Aldh1a1+ cross-correlation (Fig. 2g, top), while negative PC1 (-PC1) closely approximated the average Calb1+ or VGlut2+ cross-correlations (Fig. 2g, bottom)—which is consistent with the opposing nature of Calb1+/VGlut2+ vs Aldh1a1+ average cross-correlations (Figure 2c’-d). PC1 scores (the projection of each cross-correlation onto PC1) calculated for all recordings from each subtype revealed distributions that were mostly negative for VGlut2+ and Calb1+ axons recordings (Fig. 2h, mean = -0.46 and -0.50, p-values = 1×10^-06^ and 9×10^-05^ respectively, Wilcoxon signed-rank test), but largely positive for Aldh1a1+ axon recordings (Fig. 2h, mean = +0.26, p-value = 8×10^-06^). The PC1 score distributions were also significantly different between Aldh1a1+ vs VGlut2+ and Calb1+ (p-values = 8×10^-11^ and 1×10^-08^, respectively, Mann-Whitney U test with Bonferroni correction), but not between Calb1+ and VGlut2+ (p-value = 1) (Fig. 2i). For comparison, the same measurements from DAT-Cre labeled mice revealed a wide range of PC1 scores for axon recordings (Fig. 2h bottom, mean = -0.11) with a distribution significantly different than that of the isolated subtypes (p-values = 0.004 for VGlut2+, 0.001 for Calb1+, and 4×10^-05^ for Aldh1a1+ axons; Mann-Whitney U test with Bonferroni correction, Fig. 2i). Therefore, Calb1+ and VGlut2+ axon recordings displayed largely deceleration correlated signaling, which was markedly different from the acceleration correlated signaling of the majority of Aldh1a1+ axon recordings.

**Figure 2:**
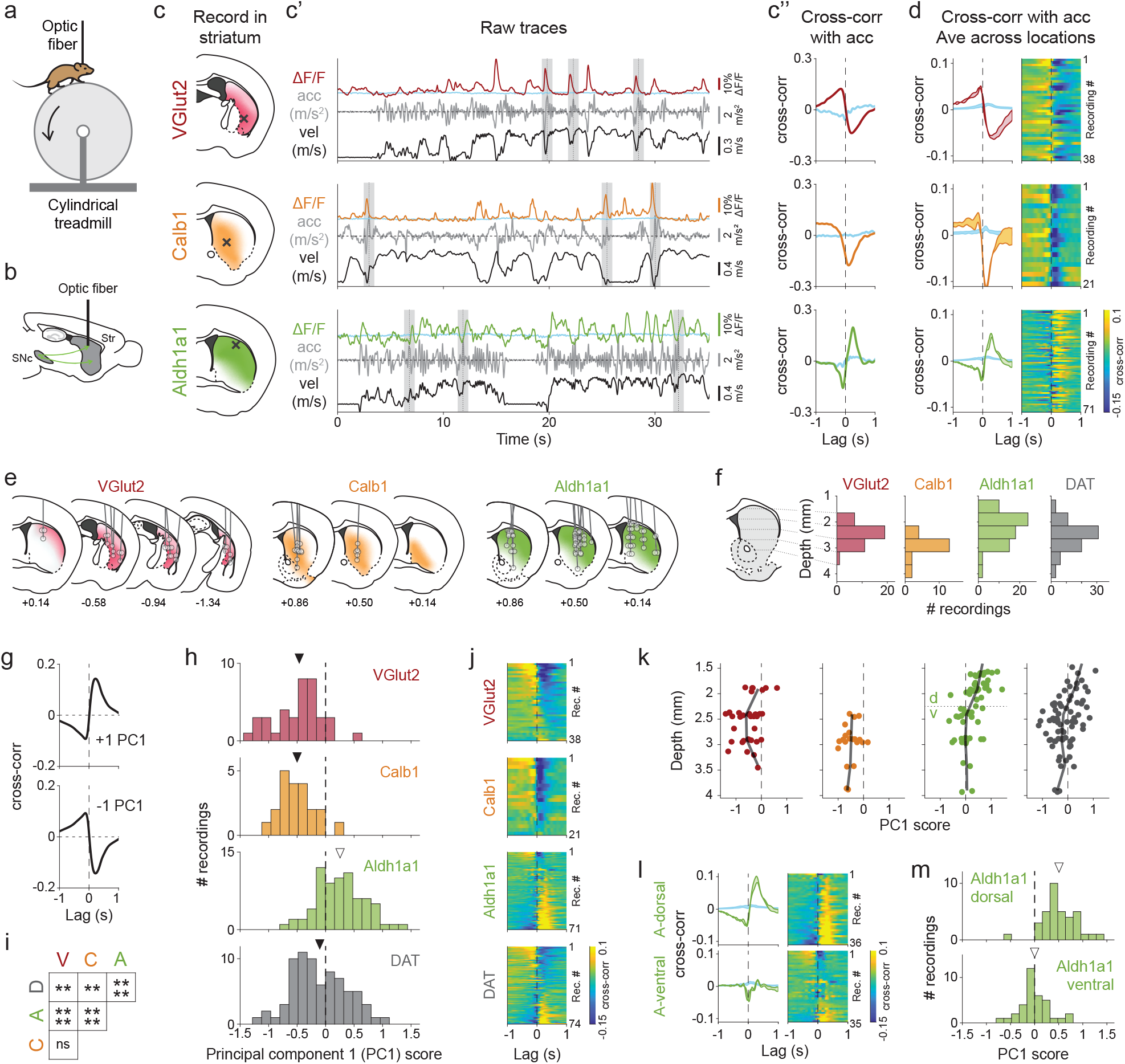
Striatal axons of genetic dopamine neuron subtypes have distinct signaling patterns during locomotion. a. Mouse running on treadmill during fiber photometry. b. Schematic of striatum recording used for Fig. 2. c. Example recordings for each subtype studied, showing recording location marked by an X (c), ΔF/F traces, acceleration and velocity (c’), and cross-correlation between ΔF/F traces and acceleration (c’’) for traces shown in c’. lsosbestic control shown in blue. Grey shaded bars in c’ show example 1s windows around large accelerations. Traces shown were recorded from mice that had never received rewards on the treadmill. d. Average cross-correlation between ΔF/F traces and acceleration for all recordings of each subtype. lsosbestic control shown in blue. Shaded regions denote mean ±. s.e.m. Heatmap shows cross-correlations for each recording, randomly sorted (VGlut2 mice = 11, n = 38 recordings; Calb1 mice = 5 recordings, n = 21; Aldh1a1 mice = 13, n = 71 recordings) e. Recording locations in striatum for recordings shown in d. Shaded colors represent projection patterns for each subtype. f. Histograms of depth from brain surface for all recordings shown in d, and recordings from DAT mice (as in Fig. 1) (0.5 mm bins) (DAT mice = 1, n = recordings). g-m. Principal component analysis (PCA) on ΔF/F-acceleration cross-correlations for all striatal recordings, including DAT. g. Representation of +PC1 shown at top and -PC1 at bottom. PC1 accounts for 63% of variance of all cross correlations. h. Distribution of principal component 1 (PC1) scores for all recordings of each subtype and DAT. Mean PC1 score for VGlut2 = -0.46, Calb1 = -0.50, Aldh1a1 = +0.26, DAT = -0.10 (means marked by arrows, ▾ = negative, ▽ = positive) i. Statistical significance between PC1 scores for all subtypes. p-values: V-D = 0.004, C-D = 0.001, A-D = 4 ×10^-05^, V-A = 8 ×10^-11^, C-A = 1 × 10^-08^, V-C = 1 (not significant), Mann-Whitney U test with Bonferroni correction. j. Cross-correlations between ΔF/F and acceleration for all recordings (same as d plus DAT) sorted by PC1 score. k. Distribution of PC1 scores along the dorso-ventral axis of the striatum. Black line represents moving average (0.5 mm bins). l. Average cross-correlations between ΔF/F and acceleration for Aldh1a1 recordings split by striatum depth (threshold = 2.25 mm, dotted line in k). lsosbestic control shown in blue. Shaded regions denote mean ± s.e.m. (A-dorsal mice = 12, n = 36 recordings; A-ventral mice = 12, n = 35 recordings). m. Distribution of PC1 scores for Aldh1a1 recordings split by depth, as in l. Mean PC1 score for A-dorsal = + 0.51, A-ventral = +0.0002 (marked by arrow as in g).

Interestingly however, while VGlut2+ and Calb1+ recordings were largely homogeneous (89% and 95% of recordings, respectively, were defined by negative PC1 scores), Aldh1a1+ recordings were more heterogeneous (70% positive and 30% negative PC1 scores). This was clearly visible when the cross-correlation heat maps were sorted by PC1 scores (Fig. 2j). This Aldh1a1 heterogeneity varied anatomically within the striatum, as shown when plotting PC1 scores vs recording depth from the brain surface for each recording (Fig. 2k): while VGlut2+ and Calb+ axon recordings were functionally homogeneous across depths (negative PC1 scores), Aldh1a1+ recordings were functionally homogeneous only in dorsal striatum (positive PC1 scores in 97% of recordings above 2.25mm from brain surface), but heterogeneous at more ventral depths (43% positive PC1 scores) (Fig. 2l-m). This matches previous reports showing that most individual axons in the dorsal-most striatum were acceleration correlated^3^, and is not unlike studies reporting functional differences in dopamine axons across a broader range of locations in striatum^3,34–37^. Therefore, while Aldh1a1+ axon recordings were overall enriched with acceleration correlations, unlike Calb1+ and VGlut2+, the recordings were not functionally homogeneous and could be further isolated in striatum through anatomical means.

Isolating genetic subtypes allowed for the identification of clear functional differences in axons; however, it is still possible that local cholinergic modulation could explain these differences, since axonal arbors of different subtypes are densest in different regions, and these regions could be differently modulated by acetylcholine (Fig. 1a). Therefore, we next asked whether the somas in SNc of the same genetic subtypes displayed similar functional signaling to that observed in their striatal axons. We recorded from a wide range of locations within SNc where somas of the three subtypes overlapped (Fig. 3c), and used the same photometry methods and behaviors (in an overlapping cohort of mice, see Supplemental Table 1) to recorded calcium transients from somas of the different subtypes (Fig. 3a-c). We found that Calb1+ and VGlut2+ somas preferentially signaled during locomotion decelerations (Fig. 3c’), exemplified by a trough at positive lags in cross-correlations between ΔF/F and acceleration (Fig. 3c”, d), similar to axons of the same subtypes. The similarity in locomotion related signaling between Calb1+ and VGlut2+ somas and axons was further tested by using the same PC1 (obtained from axonal cross-correlation PCA, Fig. 2g) to decompose somatic cross-correlations (see Methods). We found that 58% of somatic variance was explained by the axonal PC1 and, similarly to Calb1+ and VGlut2+ axons, most somatic PC1 scores were negative (Fig. 3g, mean = -0.45 for VGlut2+ and -0.54 for Calb1+). Thus, Calb1+ and VGlut2+ SNc somas displayed locomotion related signaling highly similar to their axons in striatum.

**Figure 3:**
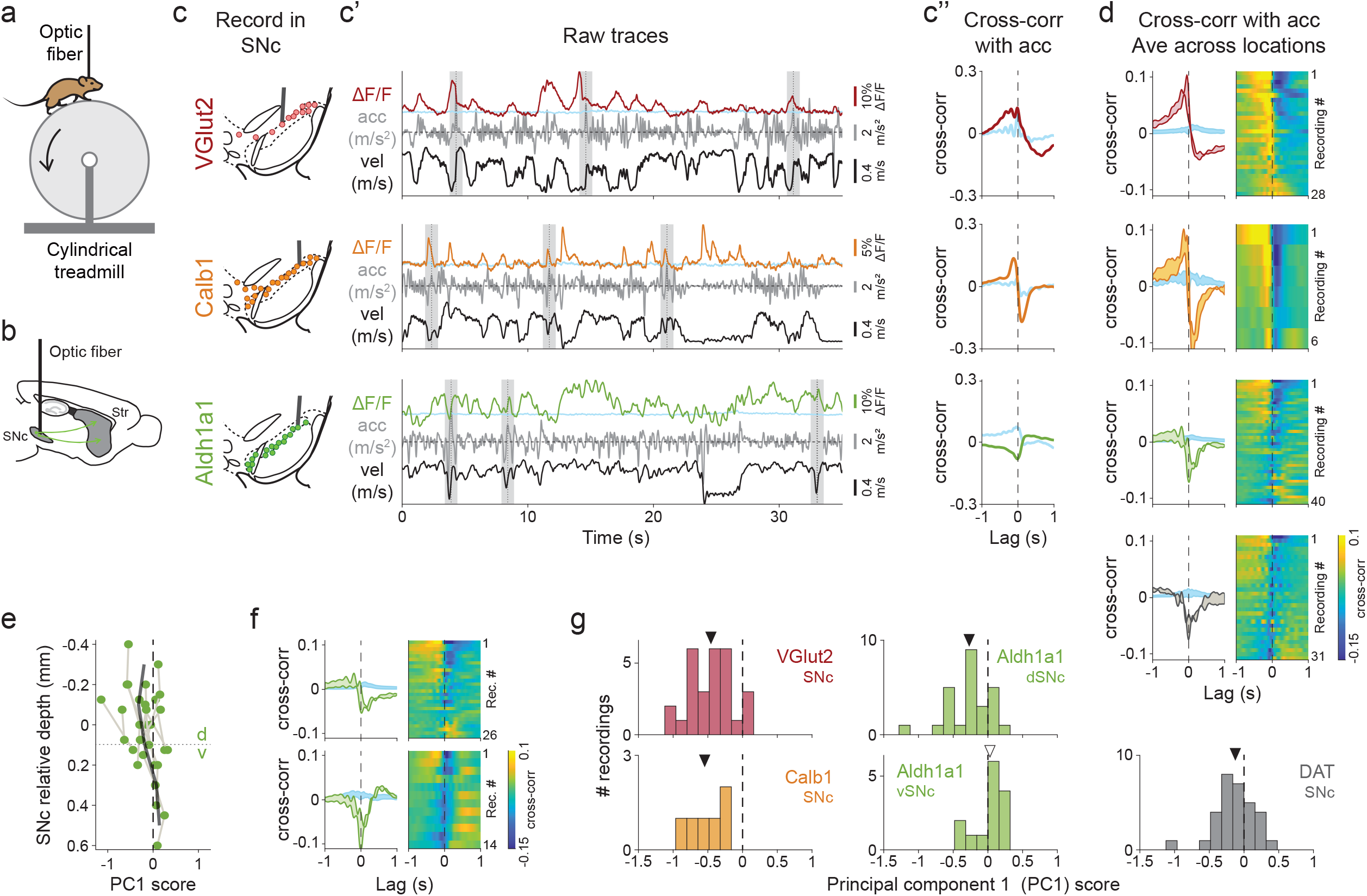
SNc somas of genetic dopamine neuron subtypes have similar signaling patterns to their axons during locomotion. a. Mouse running on treadmill during fiber photometry b. Schematic of SNc recording used for Fig. 3. c. Example recordings for each subtype studied, showing recording fiber track (c), ΔF/F traces, acceleration and velocity (c’), and-cross correlation between ΔF/F and acceleration (c’’) for traces shown in c’. lsosbestic control shown in blue. Grey shaded bars in c’ show example 1s windows around large accelerations. d. Average cross-correlation between ΔF/F traces and acceleration for all recordings of each subtype and DAT. lsosbestic control shown in blue. Shaded regions denote mean ± s.e.m. Heatmap shows cross-correlations for each recording, sorted by PC1 score (PC1 from axonal recordings, Fig. 2g). (VGlut2 mice = 11, n = 28 recordings; Calb1 mice = 3, n = 6 recordings; Aldh1a1 mice = 12, n = 40 recordings; DAT mice = 8, n = 31 recordings) e-g. Principal component analysis on ΔF/F-acceleration cross-correlations for all recordings, including DAT, using same principal component calculated in Fig. 2 for striatal recordings. This PC1 explains 56% of variance in SNc data. e. Distribution of PC1 scores along the dorso-ventral axis of the SNc for Aldh1a1 recordings. Black line represents moving average (0.2 mm bins). f. Average cross-correlation between ΔF/F traces and acceleration for Aldh1a1 recordings split by relative depth in SNc, as shown in e. lsosbestic control shown in blue. Shaded regions denote mean ± s.e.m. Heatmap shows cross-orrelations for each recording, sorted by PC1 score. (Aldh1a1-dSNc mice = 12, n = 26 recordings; Aldh1a1-vSNc mice = 6, n = 14 recordings) g. Distribution of PC1 scores for all SNc recordings of each subtype and DAT. For Aldh1a1, recordings are divided by relative depth in SNc (see e, f). Mean PC1 score for VGlut2 = -0.45, Calb1 = -0.54, Aldh1a1-dorsal = -0.27, Aldh1a1-ventral = +0.02, DAT = -0.12 (marked by arrows, ▾ = negative, ▽ = positive).

For Aldh1a1+ SNc somas we observed functional heterogeneity across recordings, similarly to their axons in striatum. While many recordings revealed preferential signaling during locomotion accelerations (Fig. 3c’-c’’), others revealed preferential signaling during decelerations (Fig. 3d). This was further exemplified by a broad spread of both positive (40%) and negative (60%) PC1 scores (same axonal PC1 from Fig. 2g) when decomposing Aldh1a1+ somatic cross-correlations (Fig. 3e,g). Importantly, compared to the 40% of Aldh1a1+ soma recordings with positive PC1 scores, only 11% of VGlut2+ and none of Calb1+ soma recordings had positive PC1 scores, and the distribution of Aldh1a1+ soma PC1 scores were significantly different from those of VGlut2+ and Calb1+ (p-values = 8×10^-04^ and 0.02 respectively, Mann-Whitney U test with Bonferroni correction). Just like the axonal recordings, Aldh1a1+ SNc recordings varied along the dorso-ventral axis. We plotted PC1 scores for Aldh1a1+ soma recordings by relative depth in SNc (see Methods), and found more negative PC1 scores in dorsal SNc Aldh1a1+ somas and more positive PC1 scores in ventral SNc somas (Fig. 3e). We identified a dorso-ventral anatomical division separating Aldh1a1+ somas into two groups with average positive vs negative PC1 scores: the ventral-Aldh1a1+ somas had significantly more positive PC1 scores vs dorsal-Aldh1a1+ somas (p-value = 0.003, Mann-Whitney U test) (Fig. 3f-g). Together with our functional analysis of axons, we conclude that the Aldh1a1+ subtype includes previously unknown functional sub-populations with different projection patterns, in accordance with reports that ventral SNc neurons project to dorsal striatum (and dorsal SNc to ventral striatum)^38^. The Aldh1a1+ sub-population with somas in ventral SNc projecting to dorsal striatum, which we here term Aldh1a1+ “Type 1”, signals preferentially during locomotion accelerations, unlike VGlut2+ and Calb1+ subtypes. Overall, using genetic labeling and anatomical distributions, we found highly similar signaling in SNc somas and striatal projection axons.

Isolating dopamine neuron subtypes therefore showed that axons and somas display similar signaling during locomotion; however, it is still possible that somas and axons could have similar correlation to movements, but low correlations to each other (for example, somas and axons could be active at different accelerations). Therefore, to provide conclusive evidence supporting the hypothesis that, within subtypes, dopaminergic axons track somatic signaling, we performed simultaneous striatal axon and SNc soma recordings. As above in DAT-Cre mice (Fig. 1b-g), we placed one optic fiber in SNc and a second fiber in striatum (Fig. 4a-b), but instead recorded from Calb1+, VGlut2+ or Aldh1a1+ Type 1 subtypes (Fig. 4c). Within subtypes, individual recordings often revealed highly similar signaling between striatal axons and SNc somas (Fig. 4c’), resulting in high cross-correlations (Fig. 4c’’), and this was consistent across recordings and mice (Fig. 4d). On average, the cross-correlation between soma and axon ΔF/F recordings was significantly higher compared to DAT+ recordings (Fig. 4d-f; mean = 0.65 for VGlut2, 0.67 for Calb1, 0.58 for Aldh1a1+ Type 1, and 0.37 for DAT+. p-values for comparison with DAT+ = 3×10^-04^ for VGlut2+,

**Figure 4:**
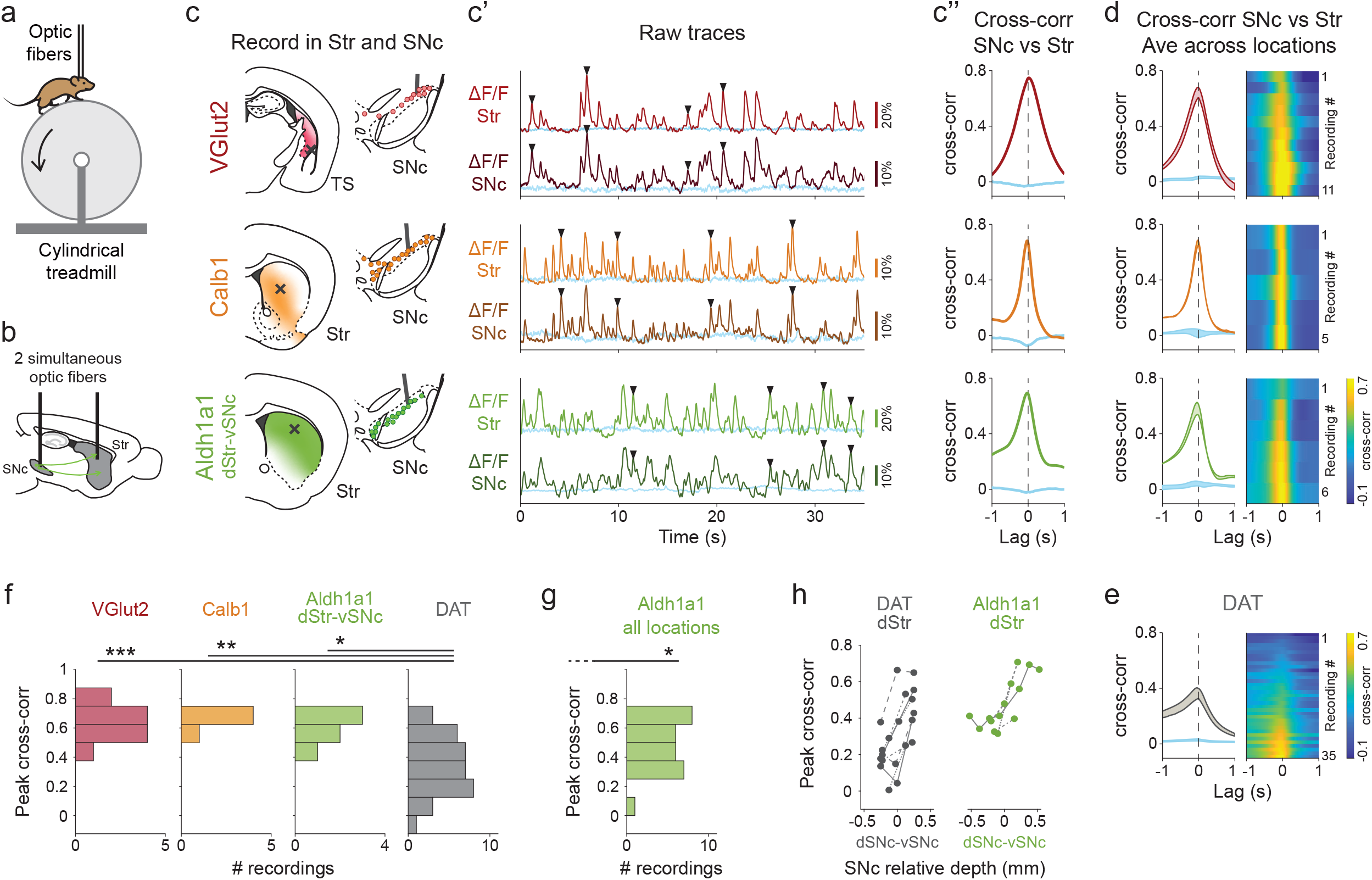
Highly correlated signaling in axons and somas within subtypes of dopamine neurons. a. Mouse running on treadmill during dual fiber photometry b. Schematic of simultaneous photometry recordings from SNc and striatum used for Fig. 4. c. Example recordings for each subtype studied, showing recording location (c), ΔF/F traces from SNc and striatum (c’), and cross correlation between simultaneous ΔF/F traces from striatum and SNc (c’’) for traces shown in c’. lsosbestic control shown in blue. ▾= example transients present in SNc and in striatum. d. Average cross correlation between simultaneous ΔF/F from striatum and SNc for all recordings of each subtype. For Aldh1a1 only pairs of recordings between dorsal striatum and ventral SNc are shown (Type 1). lsosbestic control shown in blue. Shaded regions denote mean ± s.e.m. Heatmap shows cross correlations for each paired recording sorted by peak magnitude (VGlut mice = 4, n = 11 recordings; Calb1 mice =2, n = 5 recordings; Aldh1a1-dStr-vSNc mice = 4, n = 6 recordings) e. Same as d but for A recordings (same as shown in Fig. 1f, mice = 5, n =35 recordings) f. Distribution of peak cross correlations between SNc and striatum for recordings of all subtypes and DAT (same as Fig. 1g) shown in d-e. For Aldh1a1, only pairs of recordings between dorsal-striatum and ventral-SNc are shown (Type 1).P-values for comparison to DAT: VGlut = 2×10^-04^, Calb1 = 3×10^-03^, Aldh1a1-type1 = 0.03 (Mann-Whitney U test with Bonferroni correction). g. Same as in f but including all paired recordings obtained for Aldh1a1 (mice = 8, n = 28 recordings). P-value for comparison with DAT = 0.02 (Mann-Whitney U test with Bonferroni correction). h. Peak cross correlations between dorsal striatum recordings (from Aldh1a1 or DAT) vs different relative depths in SNc.

0.003 for Calb1+, 0.03 for Aldh1a1+ Type 1, Mann-Whitney U test with Bonferroni correction). Cross-correlations between simultaneous soma and axon recordings of all Aldh1a1+ neurons (not just Type 1) were also significantly higher compared to DAT+ recordings (Fig. 4g) (mean = 0.50, vs DAT+ p-value = 0.02, Mann-Whitney U test with Bonferroni correction). Also, as expected from our anatomical functional distribution results for Aldh1a1+ axons/somas (Fig. 2k, 3e) and in confirmation of our isolation of Aldh1a1+ Type 1 neurons, we found that dorsal striatum axons were more highly correlated to ventral SNc somas than to dorsal SNc somas (Fig. 4h). Overall, we conclude that recording from isolated dopaminergic functional subtypes leads to highly similar signaling patterns between somas and axons in behaving mice.

In summary, here, we used fiber photometry to record calcium transients in both axons and somas of dopaminergic neurons in head-fixed mice running on a treadmill. We first recorded from the SNc in DAT-Cre mice, where all dopaminergic subtypes were indiscriminately labelled, and we observed large discrepancies between axonal and somatic signaling, similar to previous observations from VTA dopaminergic neurons^17^ and consistent with the idea that local cholinergic signaling plays a significant role in striatal dopamine release during behavior^16^. We then used genetic strategies to isolate three previously identified dopamine subtypes (VGlut2+, Calb1+, and Aldh1a1+), which together account for most dopamine neurons in the SNc. We observed that, in both axons and somas, VGlut2+ and Calb1+ dopamine neurons displayed deceleration correlated signaling, and that axonal and somatic signaling was highly correlated within each subtype. The third subtype, Aldh1a1+, displayed acceleration correlated signaling in both somas and axons (unlike that observed in VGlut2+ and Calb1+ neurons), though this subtype was not functionally homogeneous. Aldh1a1+ dopamine neurons could be further divided into two functional sub-populations, and when we anatomically targeted a sub-population, we again found highly correlated signaling between the axons and somas. Thus, when the diversity of dopaminergic neurons is taken into account, there is high correlation between somatic and axonal signaling, providing strong support for the classical view that striatal dopamine release is controlled by anterogradely propagating action potentials originating in midbrain somas.

Our observation of highly correlated signaling in axons and somas of dopamine neurons is also in agreement with previous reports demonstrating that cholinergic interneurons and dopamine axons in striatum are often desynchronized during behavior^27^, making it difficult to explain the majority of dopamine release based on cholinergic triggering. However, this does not exclude the possibility that local cholinergic modulation may still play a role in controlling dopamine release at specific behavioral time points. For example, striatal dopamine and acetylcholine signaling have been found to synchronize at certain times during behavior, such as at locomotion initiation or during turning^16,27^.

While the three dopaminergic subtypes studied here displayed correlations between axons and somas when isolated, our photometry recording methods average over many axons and somas, and thus it is possible that some further functional heterogeneity is contained within each subtype—as exemplified by the Aldh1a1+ subtype. This might align with further genetically defined subdivisions not yet identified^31^. Future studies will be able to investigate this question using cellular resolution^7,39^ or axonal resolution recording methods ^3^. Regardless, our results here provide strong evidence for a previously unknown connection between genetic and functional diversity in dopamine neurons and show that axons track somatic signaling within dopaminergic subtypes, which must be considered in order to fully understand the mechanisms of dopamine release in striatum during behavior.

## Methods

### RESOURCE AVAILABILITY

#### Lead contact

Further information and requests for resources and reagents should be directed to and will be fulfilled by the lead contacts, Daniel A. Dombeck and Rajeshwar Awatramani.

#### Materials availability

Mouse lines generated in this study will be shared upon request upon completion of Material Transfer Agreement due to institutional policy and will be deposited to a mouse repository (e.g., MMRRC).

#### Data and code availability

All data has been deposited at GitHub and is publicly available as of the date of publication. DOIs are listed in the key resources table.

All original code has been deposited at GitHub and is publicly available as of the date of publication. The link to the GitHub repository is included in the key resources table.

Any additional information required to reanalyze the data reported in this paper is available from the lead contact upon request.

## EXPERIMENTAL MODEL AND SUBJECT DETAILS

### Mice

All animals used in this study were maintained and cared following protocols approved by the Northwestern Animal Care and Use Committee. Cre mouse lines were maintained heterozygous by breeding to wild-type C57BL6 mice. The Th-Flpo line and the Ai93D reporter line were maintained homozygous. The Dat-tTA mouse line was maintained heterozygous by breeding with the Ai93D reporter. The Aldh1a1-iCre line was generated at Northwestern University by the Transgenic and Targeted Mutagenesis Laboratory. Mice were genotyped using primers detailed in the Key resources table.

Both males and females were used for all experiments. Adult mice were used for viral injections at 2 to 4 months old. For indiscrimitate labelling of SNc dopamine neurons, DAT-IRES-Cre mice (RRID:IMSR_JAX:027178) were injected with AAV1-CAG-FLEX-GCaMP6f virus (RRID:Addgene_100835). For labelling VGlut2+ or Aldh1a1+ dopamine neurons, we crossed VGlut2-IRES-Cre (RRID:IMSR_JAX:016963) or Aldh1a1-iCre mice (new line) with Th-2A-Flpo mice^23^, and offspring were injected with AAV8-EF1α-CreOn/FlpOn-GCaMP6f virus (RRID:Addgene_137122). For labelling Calb1+ dopamine neurons, we crossed Calb1-IRES2-Cre mice (RRID:IMSR_JAX:028532) with DAT-tTA (RRID:IMSR_JAX:027178), Ai93D (CreOn/tTAOn GCAMP6f reporter) (RRID:IMSR_JAX:024107) mice.

## METHOD DETAILS

### Generation and characterization of the Aldh1a1-iCre line

Because our previous Aldh1a1-CreERT2 strain displayed substantial mosaicism, resulting in only weak GCaMP6f signals, we opted to generate an Aldh1a1-iCre strain. The Aldh1a1-iCre line was generated at Northwestern University by the Transgenic and Targeted Mutagenesis Laboratory. In brief, a P2A peptide directly followed by iCre and a BGH polyA sequence were inserted after the last encoded amino acid of Aldh1a1, using CRISPR mediated HDR (Guides 1-2 – see Key resources table). First, PRXB6/N ES cells were electroporated and screened for insertion and correct locus with multiple primer pairs (Aldh1a1-iCre insertion primers Forward 1-3 and Reverse 1-3 – see Key resources table) followed by Sanger sequencing of iCre+ clones from outside the homology arms through the construct in order to confirm fidelity of the insertion. Clone C7 was expanded and injected into blastocysts to generate chimeras and used for all experiments herein.

Aldh1a1-iCre mice were genotyped using primer set 3 described above. To determine the expression fidelity of this allele, 0.4 µl of AAV5-EF1α-DIO-mCherry (RRID:Addgene_37083) was injected into SNc bilaterally (coordinates relative to bregma: x = ±1.45mm, y = -3.15mm, z = -3.1, -4.1, -4.4, -4.7mm, 0.1 µl at each depth) in n = 4 adult mice. Three weeks later, mice were perfused, and brains were sectioned at 25 µm for immunofluorescence staining. Floating sections were first blocked for 24 hours at 4°C in PBS containing 0.03% Triton-X and 5% normal donkey serum. Sections were incubated with primary antibodies against Aldh1a1 (goat, R&D Systems Cat# AF5869, RRID:AB_2044597), TH (mouse, Sigma-Aldrich Cat# T2928, RRID:AB_477569 ; Pel-Freez Biologicals Cat# P40101-0, RRID:AB_461064) and mCherry (rat, Thermo Fisher Scientific Cat# M11217, RRID:AB_2536611) in blocking buffer for 24 hours, followed by 4 washes in PBS-Tween20 and incubation with secondary antibodies (Donkey anti Goat Alexa Fluor 488 [Molecular Probes Cat# A-11055, RRID:AB_2534102], Donkey anti Mouse Alexa Fluor 647 [Thermo Fisher Scientific Cat# A-31571, RRID:AB_162542], Donkey anti Rabbit Alexa Fluor 647 [Thermo Fisher Scientific Cat# A-31573, RRID:AB_2536183], Donkey anti Rat Cy3 [Jackson ImmunoResearch Labs Cat# 712-165-153, RRID:AB_2340667], and DAPI [Thermo Scientific 62248]) for 2 hours at room temperature. Sections were then imaged at 20x. For each brain, 4-5 sections spaced at least 100 microns apart and centered about the area of maximal viral recombination were counted for mCherry+/DAPI+/Aldh1a1+ and mCherry+/DAPI+/Aldh1a1-cells (2740 cells total).

### Stereotaxic viral injections

Adult mice (postnatal 2-4 months old) were anesthetized with isoflurane (1-2%), and a 0.5–1-mm diameter craniotomy was made over the right substantia nigra (-3.25 mm caudal, +1.55 lateral from bregma). A small volume (0.4 μl total) of virus (AAV8-EF1α-CreOn/FlpOn-GCaMP6f (RRID:Addgene_137122, titer 6.10E+13) for Aldh1a1-iCre/Th-Flpo and VGlut2-IRES-Cre/Th-Flpo mice, or AAV1-CAG-FLEX-GCaMP6f (RRID:Addgene_100835, titer 2.00E+13) for DAT-Cre mice), diluted 1:1 in PBS, was pressure injected through a pulled glass micropipette into the SNc at 4 depths (-3.8, -4.1, -4.4 and -4.7 mm ventral from dura surface, 0.1 μl per depth). Following the injections, the skull and craniotomy were sealed with Metabond (Parkell) and a custom metal headplate was installed for head fixation. The location of recording sites was marked the surface of the Metabond for future access. For Calb1-IRES2-Cre/DAT-tTA/Ai93D mice, which express GCaMP6f endogenously, no injection was conducted and only the headplate was implanted at this time. 4 weeks were allowed for GCaMP6f expression to ramp up and fill dopaminergic somas in SNc and axons in striatum.

### Training and behavior

Starting 1-2 weeks after injection, mice were head-fixed with their limbs resting on a 1D cylindrical Styrofoam treadmill ∼20 cm in diameter by 13 cm wide in the dark. Mice were habituated on the treadmill for 3-10 days until they ran freely and spontaneously transitioned between resting and running. Rotational velocity of the treadmill during locomotion was sampled at 1,000 Hz by a rotary encoder (E2-5000, US Digital) attached to the axel of the treadmill and a custom LabView program.

After mice ran freely, a subset of mice were water restricted and received unexpected water rewards, aversive air puffs, and light stimuli while on the treadmill, using a custom LabView program (see Supplemental Table 1 for list of which sessions/mice received these stimuli). We did not observe significant differences in cross-correlations between SNc and striatum ΔF/F during these discrete events compared to the locomotion and rest periods (data not shown); locomotion and rest periods accounted for most of the time during recording sessions. Large (16 μl) and small volume (4 μl) water rewards were delivered through a waterspout gated electronically through a solenoid valve, which was accompanied by a short ‘click’ noise. Air puffs were delivered by a small spout pointed at their left whiskers, which was connected to a ∼20 psi compressed air source and triggered electronically through the opening of a solenoid valve for 0.2s. Triggering of this solenoid was also accompanied by a ‘click’ noise. For light stimuli, a blue LED placed ∼30 cm in front of the head-fixed mouse was electronically triggered for 0.2s. Rewards, air puffs and light stimuli were alternated at random during recordings and delivered at pseudo-random time intervals (10-30s between any two stimuli).

### Fiber photometry

4 weeks after injection, mice were once again anesthetized, and a small craniotomy (1 mm in diameter) was drilled through the Metabond and skull, leaving the dura and cortex intact. Craniotomies were made at different locations depending on the experiment, which were pre-marked during the injection surgery: for SNc -3.25 mm caudal, +1.55 lateral from bregma, and different locations over striatum (for example -1.1 mm caudal +2.8 lateral, or +0.5 mm caudal + 1.8 lateral). The craniotomies were then sealed with Kwik-Sil (World Precision Instruments KWIK-SIL).

After the mice recovered from this short (10-15 min) surgery for one day, they were head-fixed on the linear treadmill, and the Kwik-Sil covering the craniotomies was removed. One or two optical fibers (200 μm diameter, 0.57 NA, Doric MFP_200/230/900-0.57_1.5m_FC-FLT_LAF) were lowered slowly (5 μm/s) using a micromanipulator (Sutter MP285) into the brain to various depths measured from the dura surface. In the striatum, recording depths ranged from 1.6 to 4.1 mm; in SNc, depths ranged from 3.5 to 4.5 mm. Recordings started at 1.6 mm in striatum, and 3.5 mm in SNc, but if no ΔF/F transients were detected at those depths the fiber was moved down in increments of 0.25-0.5 mm in striatum or 0.15-0.2 mm in SNc, until transients were detected. From there, a 15 min recording was obtained, and the fiber was moved further down in the same increments. Subsequent recordings were obtained until a depth was reached where transients were no longer detected, at which point the fiber was pulled out of the brain slowly (5 μm/s).

A custom-made photometry setup was used for recording. Blue excitation (470 nm LED, Thor Labs M70F3) and purple excitation light (for the isosbestic control) (405 nm LED, Thor Labs M405FP1) were coupled into the optic fiber such that a power of 0.75 mW emanated from the fiber tip. 470 and 405 nm excitation was alternated at 100 Hz using a waveform generator, each filtered with a corresponding filter (Semrock FF01-406/15-25 and Semrock FF02-472/30-25) and combined with a dichroic mirror (Chroma Tech Corp T425lpxr). Green fluorescence was separated from the excitation light by a dichroic mirror (Chroma Tech Corp T505lpxr) and further filtered (Semrock FF01-540/50-25) before collection using a GaAsP PMT (H10770PA-40, Hamamatsu; signal amplified using Stanford Research Systems SR570 preamplifier). A Picoscope data acquisition system was used to record and synchronize fluorescence and treadmill velocity at a sampling rate of 4 kHz.

### Histology

Immediately after the last recording, mice were perfused transcardially with PBS (Fisher) then 4% paraformaldehyde (EMS). Brains were stored in PFA at 4 °C overnight then transferred to 40% sucrose (Sigma) for at least 2 days before sectioning. Coronal slices (50 μm thick) were cut on a freezing microtome and stored at 4 °C in PBS. For immunostaining of dopaminergic neurons, sections were washed in PBS, blocked in PBS + 0.3% Triton-X (Sigma) + 5% normal donkey serum (Sigma), incubated overnight with primary antibodies Sheep anti-Tyrosine Hydroxylase (1:1000 dilution, RRID:AB_461070) and Rabbit anti-GFP, which recognizes GCaMP6f (1:1000 dilution, RRID:AB_221569), washed again in PBS + 0.3% Triton-X, then incubated with secondary antibodies tagging Tyrosine Hydroxylase with Alexa Fluor 555 (Donkey anti-Sheep Alexa Fluor 555, RRID:AB_2535857) and GCaMP6f with Alexa Fluor 488 (Donkey anti-Rabbit Alexa Fluor 488, RRID:AB_2313584). Images of SNc and Str were acquired on an Olympus or Keyence Slide Scanner (VS120 or BZ-X810, respectively) for verification of injection accuracy and fiber placement. Other brains were mounted and imaged without immunostaining for fiber placement. Histology is not available for 2 out of 12 DAT mice and 1 out of 5 Calb1 mice, and fiber tracks could not be identified for 2 out of 16 VGlut2 mice.

## QUANTIFICATION AND STATISTICAL ANALYSIS

Data was analyzed using custom MATLAB code.

### Data preprocessing

Simultaneous traces (velocity from rotary encoder, fluorescence detected by PMTs from one or two optic fibers, and output from waveform generator used to alternate 405 and 470 nm illumination every 10 ms) were collected at 4 kHz by a Picoscope data acquisition system. Fluorescence collected during 405 or 470 nm illumination (20 time bins for each pulse of 405 or 470 nm excitation) was separated using the binary output from the waveform generator. For each transition period between illumination sources, 5 time bins were excluded to remove transition times. Traces were then re-binned to 100 Hz by averaging every 40 time bins for velocity and every 40 time bins for 405 and 470 fluorescents (but only including 15 of 40 bins for each source: excluding 20 bins when the alternate source was on and 5 transition bins).

We first corrected fluorescence traces for background signal (intrinsic fluorescence and any illumination bleed-through) by subtracting 85% of the baseline (baseline defined as 8^th^ percentile over a 20s window). This 85% was estimated from photometry recordings from cortex, which was unlabeled (no GCaMP expression), obtained from 10 recordings from 5 mice. 405 and 470 fluorescence traces were corrected independently. To calculate ΔF/F, traces were then normalized by baseline fluorescence division (8^th^ percentile over a 20s window) separately for 405 and 470. The subtraction and normalization steps together corrected for bleaching and removed any slow drifts in baseline. Next, traces were converted to ΔF/F units (baseline at 0) by subtracting the baseline (median of all non-transient bins for 470 nm traces, and median of all bins for 405 nm traces)

For comparison of traces between dopaminergic subtypes, ΔF/F traces were normalized so that the baseline remained at 0 and the largest transient peak for each trace was 1. 405 traces were normalized using the amplitude of the largest peak from the corresponding 470 traces.

### Criteria for recording inclusion

Only recordings with signal-to-noise ratios greater than 10 were included in the analysis. To calculate signal-to-noise ratios for each recording, we selected well-isolated transients, as defined by having a large, fast rise (30 ΔF/F/s) immediately followed by a decay. We first removed all slow fluctuations except transients in (non-normalized) ΔF/F traces by subtracting the 8^th^ percentile over a window 2-3 times the width of observed ΔF/F transients (250 bins, 2.5s), and then smoothed the resulting trace over a 0.2s window (20 bins) to reduce noise. Transient rises and decays were identified by locating the zero-crossings on the derivative of the trace, also smoothed over 0.2s window. Only clearly isolated transients were included – those with a rise greater than 30 ΔF/F/s followed by a decay greater than -5 ΔF/F/s. Traces with less than 0.2 transients per second were excluded. Signal values for each recording were calculated as the 80^th^ percentile of isolated transient peaks. Noise for each recording was calculated by smoothing each (non-normalized) ΔF/F trace over 10 bins (0.1s), then subtracting this smoothed trace from the original ΔF/F trace and using the standard deviation of the resulting trace as the noise value. The signal and noise values were divided to obtain signal-to-noise for each trace. These steps for determining signaling to noise for each trace were not used for any further analysis.

ΔF/F traces from 405 nm illumination (isosbestic control) were used to remove any movement artifacts. While GCaMP6f fluorescence intensity is dependent on calcium concentration when excited with 470 nm light, it is still fluorescent but in a calcium-independent way when excited with 405 nm light^26^. Therefore, calcium transients in neurons are detected with 470 nm illumination but are absent with 405 nm illumination, while movement artifacts are present in both traces. Movement artifacts were identified using the 405 nm traces from each recording as follows. (Non-normalized) 405 ΔF/F traces were smoothed over a 10-bin window (0.1s). This smoothed trace was subtracted from the original 405 ΔF/F trace, so that only the noise remained (same process as used above for 470 traces to separate noise and signal). A max noise value was calculated as the max absolute value of this noise trace. Any bins in the original 405 ΔF/F trace more than 3 times this max noise (or 3 times below -max noise) were excluded from further analysis. Additionally, any sequential bins that were above max noise (or below -max noise) for longer than 0.2 s (20 bins, less than half the width of observed calcium transients) were also excluded, with an additional 0.1s (10 bins) on both sides also excluded. Any bins removed from the 405 ΔF/F trace were also removed in the corresponding 470 ΔF/F and velocity traces. If more than 5% of the bins in a recording met these movement artifact exclusion criteria, the entire recording was excluded.

### Cross-correlation between ΔF/F and acceleration

Only locomotion time bins were included for cross-correlation analysis in Fig. 2, 3. Locomotion vs rest bins were selected using a double threshold on the velocity trace in both positive and negative directions (thresh1 = ± 0.024 m/s, thresh2 = ± 0.010 m/s). Isolated 1 bin-long locomotion periods (no other movement within 2 bins on either side) were excluded, as well as rest periods shorter than 0.5 s. Time bins were considered as locomotion periods only if they lasted longer than 0.5 s and had an average velocity greater than 0.2 m/s. For a recording to be included in the cross-correlation (between ΔF/F and acceleration) analysis in Fig. 2 and Fig. 3, the recording needed to include a total of at least 100 sec of locomotion.

Acceleration was calculated from the velocity traces as the difference between consecutive treadmill velocity time bins (first smoothed over 6 bins, 0.06s), then multiplied by the sampling frequency (100 Hz) for proper m/s^2^ units. Cross-correlations between ΔF/F and acceleration were calculated for locomotion periods only (bins define above) using MATLAB’s *crosscorr* function over a 1s lag window (100 time bins). The same process was used to calculate the cross-correlation between corresponding 405 ΔF/F traces and acceleration, and any recording with a peak cross-correlation (between 405 ΔF/F trace and acceleration) above 0.1 was excluded.

For plotting in Fig. 2c,d,l and Fig. 3c,d,f, 405 and 470 cross-correlations were smoothed over 5 time-lag bins (0.05s). Shaded areas in Fig. 2d,l and 3d,f represent the mean ± standard error of the mean (s.e.m.), while accompanying heatmaps show cross-correlations for all recordings.

### Principal component analysis (PCA)

Principal component analysis (PCA) was applied to the matrix of all cross-correlation traces from striatal recordings (shown in Fig. 2d, right), from all subtypes (VGlut2, Calb1, Aldh1a1, and DAT), using MATLAB’s *pca* function (without centering: *‘Centered’, ‘off’*; however, equivalent results were obtained when we repeated the PCA analysis with centering, data not shown). This function outputs the principal components (loadings, eigenvectors), such as the first principal component, PC1 (shown in Fig. 2g), the scores for each recording’s cross-correlation along each principal component (matrix of all SNc cross-correlation traces multiplied by the loadings matrix), and the variance explained by each principal component across all recordings.

The cross-correlations for all SNc recordings (between ΔF/F and acceleration), shown in Fig. 3d right, were decomposed using the same principal components calculated above from the striatal cross correlations. Scores for SNc cross-correlations were calculated by multiplying the matrix of all SNc cross-correlation traces by the striatal loadings matrix (principal components). The % of SNc variance explained by each principal component was calculated as the variance without the mean subtracted (not centered).

P-values for reporting statistical significance for each subtype’s deviation from PC1 score = 0 (in the main text) used a non-parametric statistical test (Wilcoxon signed-rank test). For comparison of PC1 scores between each subtype and DAT or between different subtypes (Fig. 2i, 2m, 3g) we used a non-parametric statistical test for two independent populations (Mann-Whitney U test, also called Wilcoxon rank-sum test), with Bonferroni correction (p-values were multiplied by the number of comparisons performed).

### Cross-correlation between SNc and striatum ΔF/F traces

All simultaneously recorded pairs of SNc/striatum recordings where both traces had a signal-to-noise ratio above 10 were included (see above), regardless of locomotion behavior. Cross-correlations between SNc and striatum ΔF/F traces were calculated using MATLAB’s *crosscorr* function over a 1s time-lag window (100 bins). For the isosbestic control cross-correlation shown in Fig. 1e,f and Fig. 4c’’,d,e, we calculated the cross-correlations between SNc-470 and striatum-405 ΔF/F traces and also between SNc-405 and striatum-470 ΔF/F traces, and averaged the resulting cross-correlation traces together. Any pairs of recordings with a peak 405/470 average cross-correlation above 0.12 were excluded.

For plotting in Fig. 1e,f and Fig. 4c’’,d,e, 405 and 470 cross-correlations were smoothed over 5 bins (0.05s). Shaded areas in Fig. 1f and 4d,e represent the mean ± standard error of the mean (s.e.m.), while accompanying heatmaps show cross-correlations for all recordings. For comparison of peak cross correlations between each subtype and DAT (Fig. 4f. 4g) we used a non-parametric statistical test for two independent populations (Mann-Whitney U test, also called Wilcoxon rank-sum test), with Bonferroni correction (p-values were multiplied by the number of comparisons performed).

### Fiber placement localization and depth calculation

For striatum recordings, depth from surface as shown in Fig. 2f, 2k was defined as the depth at which the fiber tip was located from the brain surface, as measured by the micromanipulator used to move the fiber during photometry. To reduce overlap between data points at the same depth plotted in Fig. 2k, a random amount between + 0.05 and -0.05 mm was added to each depth. For SNc recordings, due to the tilted nature of its main axis with respect to the brain surface and due to its small width (in depth), we used a measure of relative depth (Fig. 3e). To calculate this, for each session we considered dorsal-SNc the most dorsal depth at which significant transients were observed, and ventral-SNc the most ventral depth at which transients were observed. Relative depth was then measured with respect to the middle between the dorsal and ventral most depths, in mm. To reduce overlap between data points at the same depth plotted in Fig. 3e, a random amount between +0.05 and -0.05 mm was added to each depth.

For the representation of recording locations in striatum shown in Fig. 2e (also Fig. 2c, 4c), 20x magnification images of striatum were acquired on a Keyence slide scanner (BZ-X810) (see Method details, Histology). For the slice in each brain with the clearest fiber track, fiber tracks were marked onto the images. We then identified the closest reference slice for each imaged brain slice (reference slices from the Paxinos Mouse brain atlas), spaced 0.36 mm (bregma +0.86, +0.50, +0.14, -0.22, -0.58, -0.94, -1.34 mm; as shown in schematics in Fig. 2e), and uniformly scaled this reference to approximately match the imaged slice. Recording locations (only for recordings included in Fig. 2) for each mouse were then marked on each slice, measuring depth from brain surface along the fiber track as described above. The fiber tracks and recording locations mapped to these reference slides from all mice for each subtype were combined for Fig. 2e. Circles represent approximate light collection recording area for our 200 µm fibers (∼300 µm in diameter). For representations of SNc recording locations in Fig. 3c, 4c, fiber tracks were marked onto the slices with the clearest fiber track, and the closest reference slice from the Paxinos Mouse brain atlas (bregma -2.92, -3.16, -3.40, -3.64) was identified.

## KEY RESOURCES TABLE

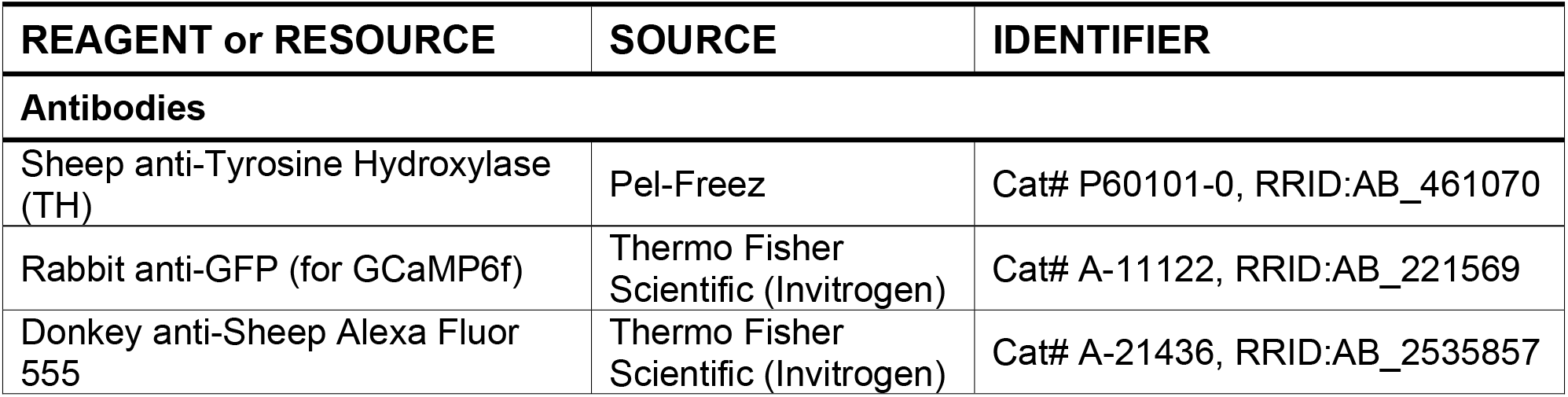
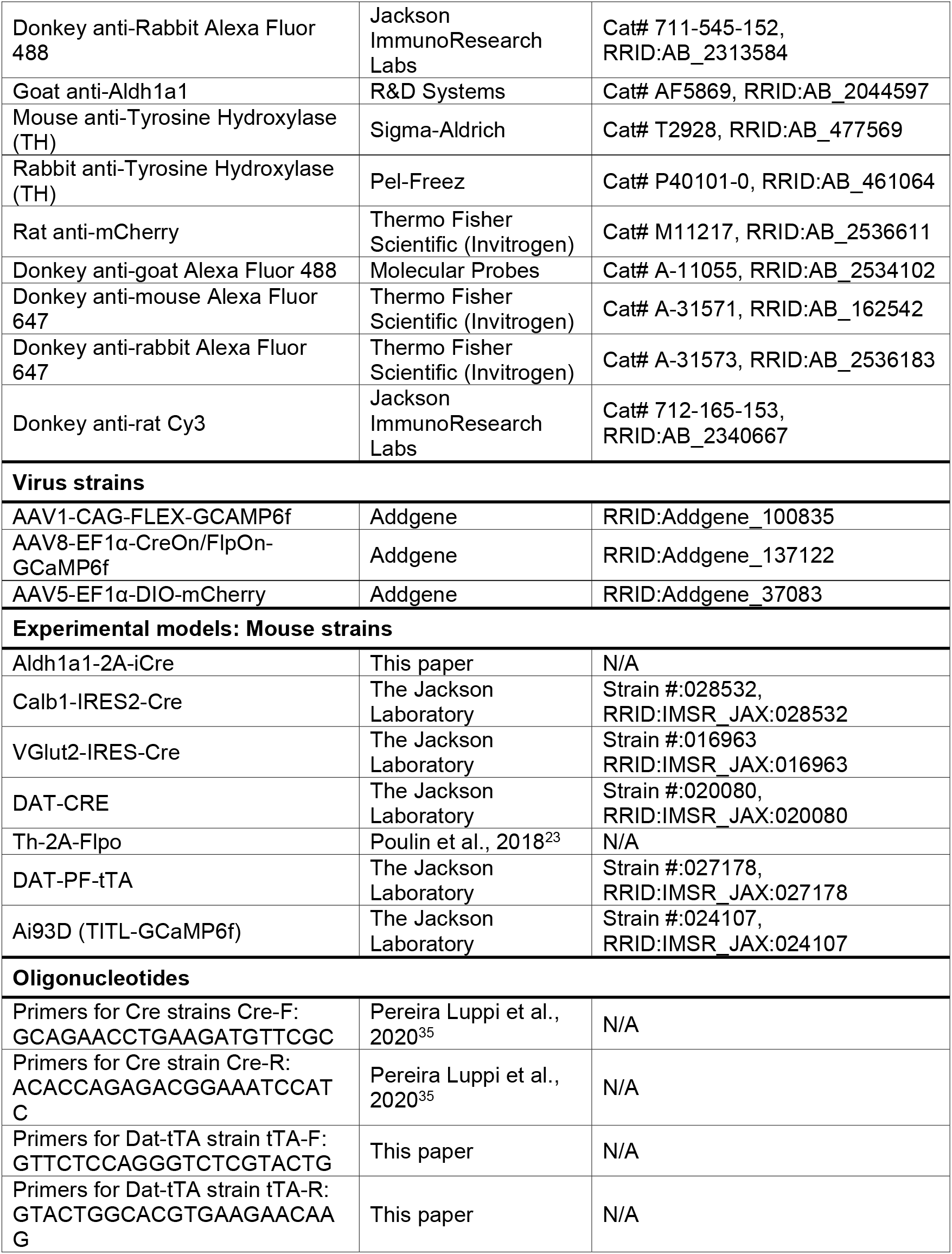
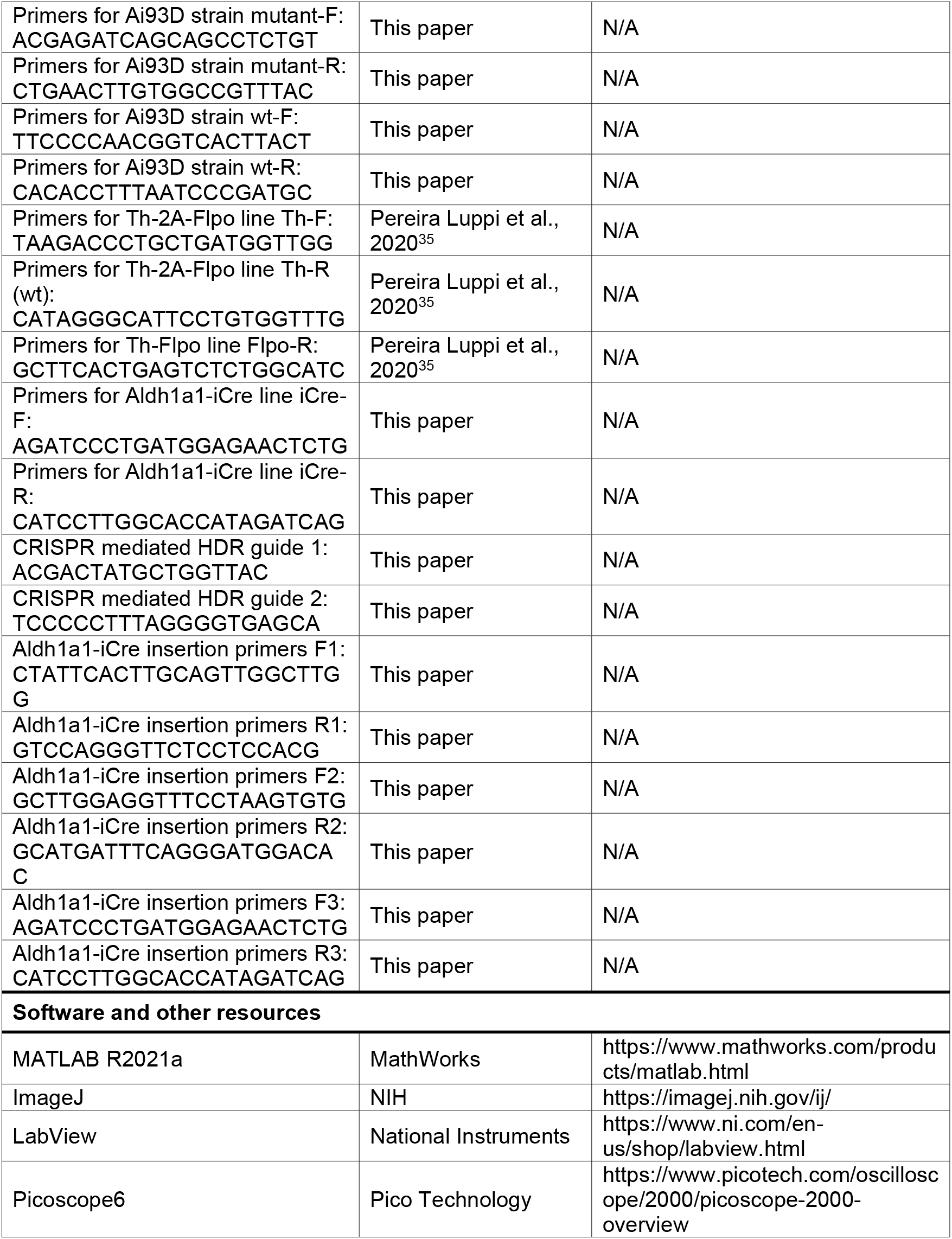
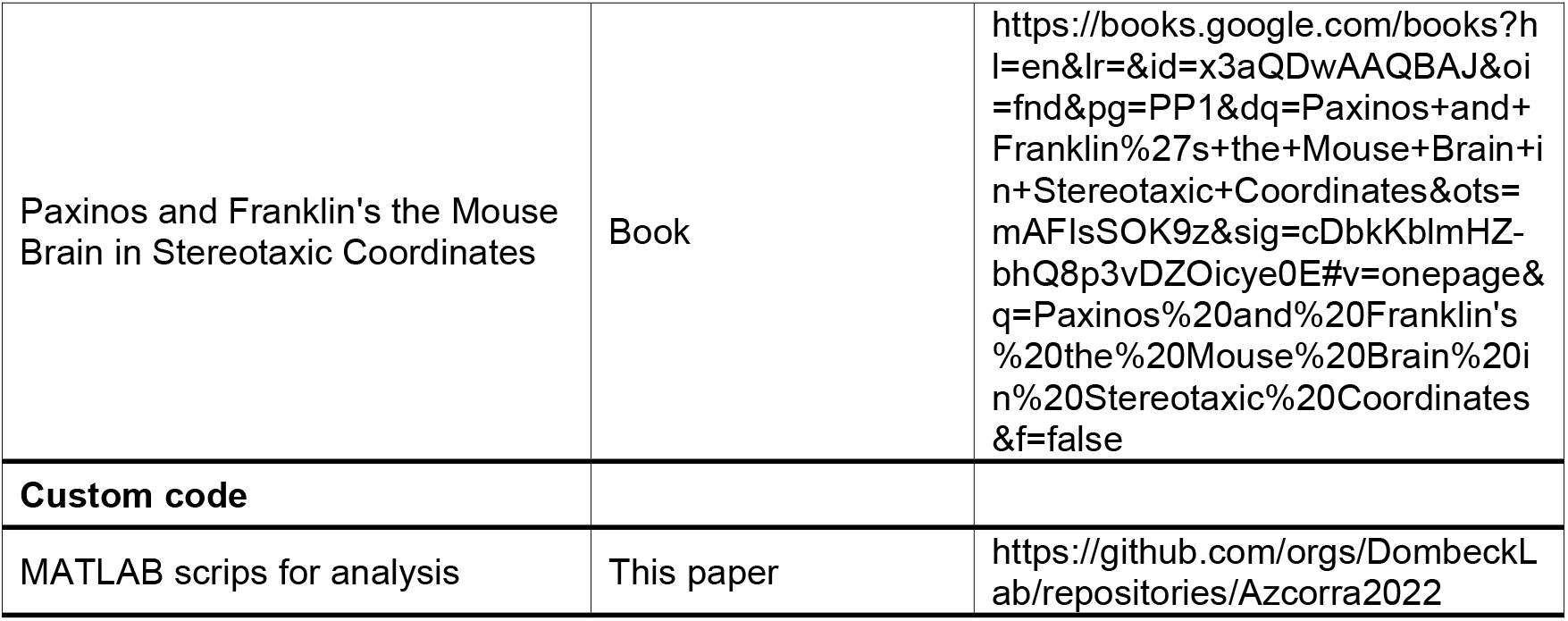

**Supplemental Figure 1:**
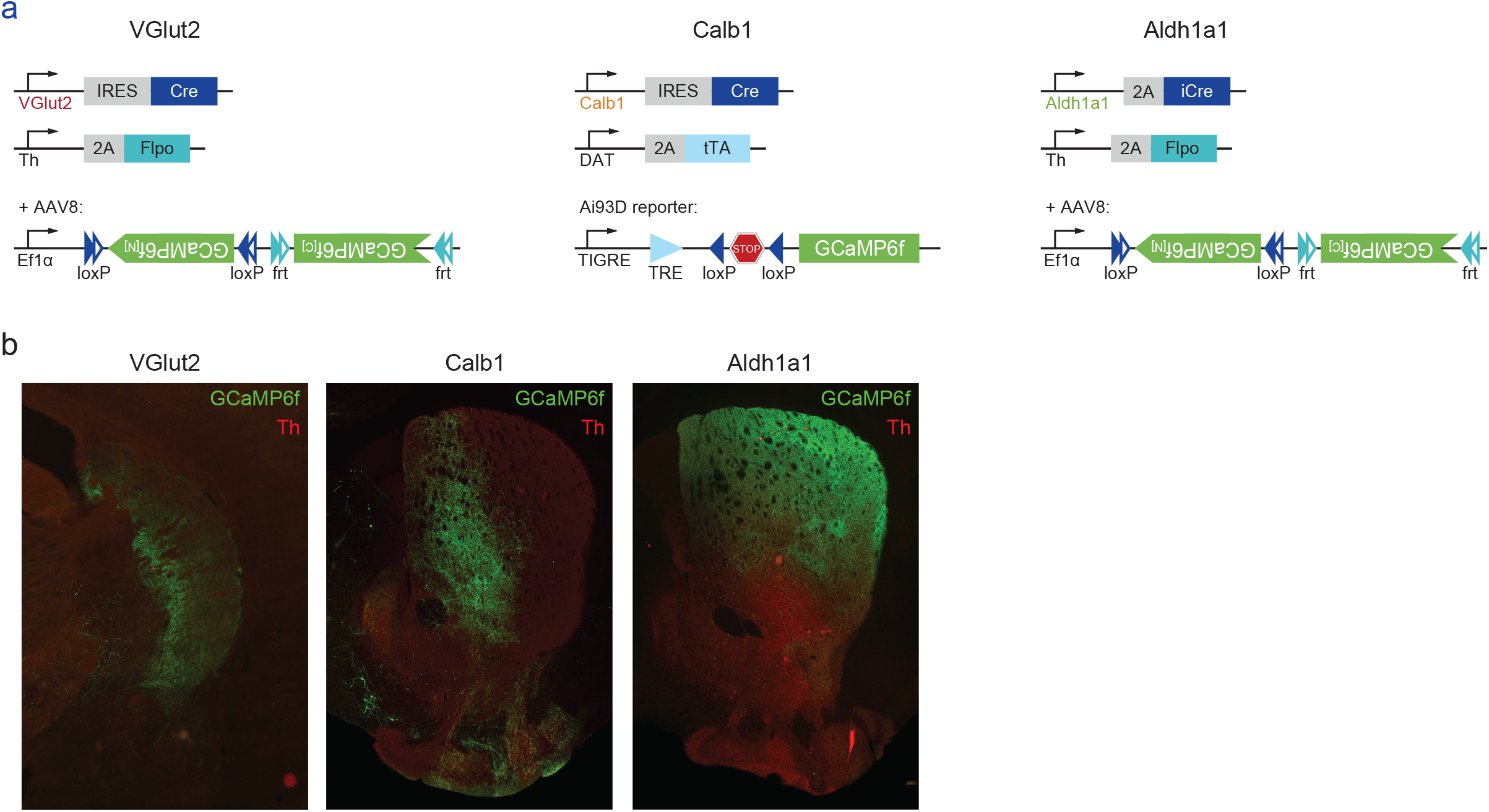
Labelling of subtypes and projection patterns. a. Intersectional genetic strategy used to label with GCaMP6f each of the subtypes studied. b. Representative slices showing axonal projections of each of the subtypes studied in striatum.

**Supplemental Figure 2:**
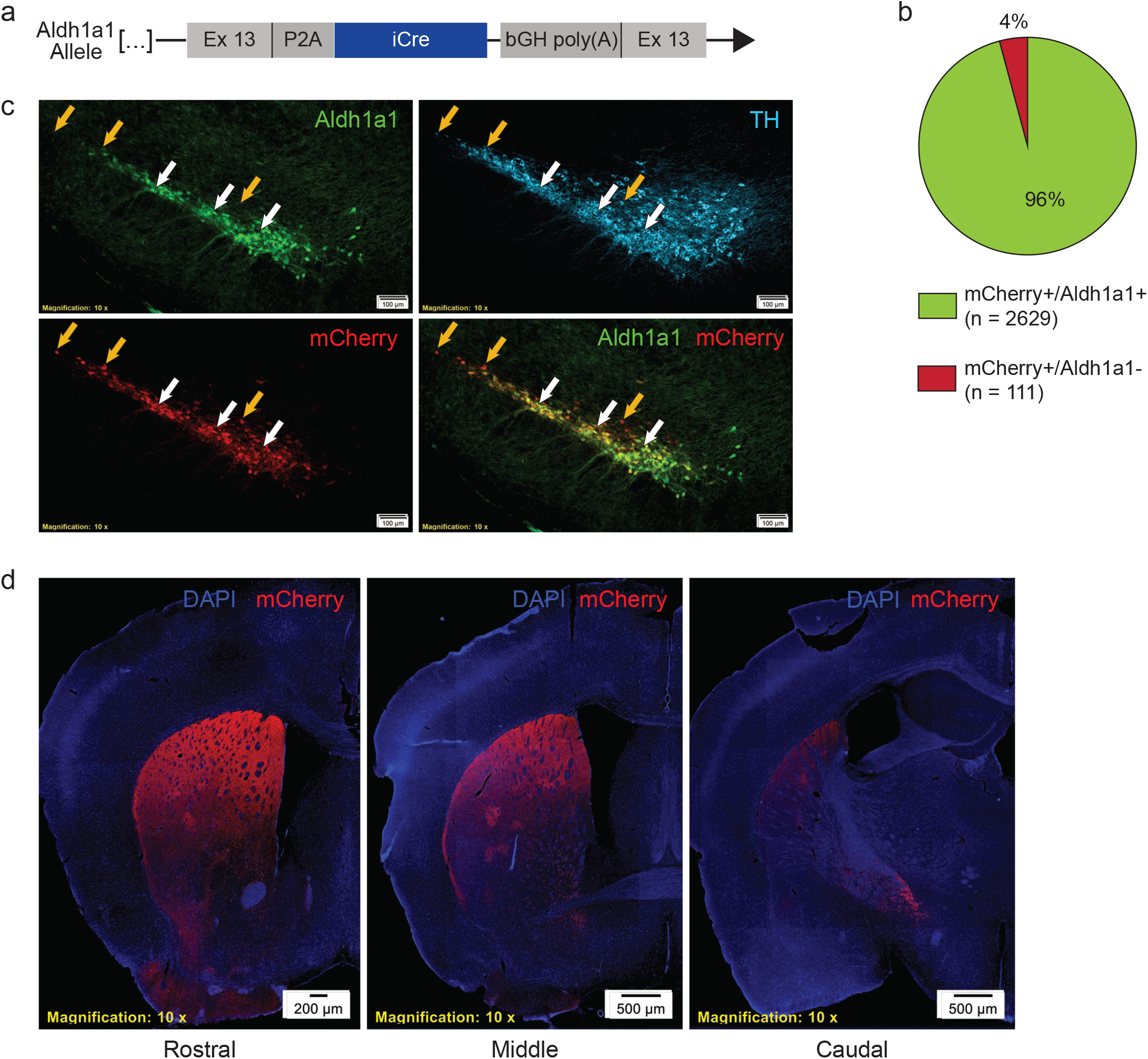
Validation of Aldh1a1-iCre mouse line. a. Schematic representation of Aldh1a1-iCre transgenic line. Endogenous Aldh1a1 gene was targeted for insertion of a P2A peptide and iCre immediately following the peptide encoded by Exon 13. b. Ratio of mCherry virally labelled cells co-staining for Aldh1a1 (n=4 mice). c. Substantia nigra pars compacta immunofluorescence staining from Aldh1a1-iCre mice injected with an AAV5-DIO-mCherry virus. Co-staining shows excellent efficiency and fidelity of iCre recombination, which is notably limited to TH+ cells in this region. Orange arrows: examples of mCherry and Aldh1a1 co-stained cells. White arrows: mCherry-expressing cells with undetectable Aldh1a1 staining, which were primarily localized to the dorsal and lateral SNc. d. Striatal sections of injected mice displaying the projection pattern of Aldh1a1-expressing DA neurons. These projections were enriched dorsally and rostrally, with significant enrichment in striatal patch compartments relative to matrix.

**Supplemental Table 1.**
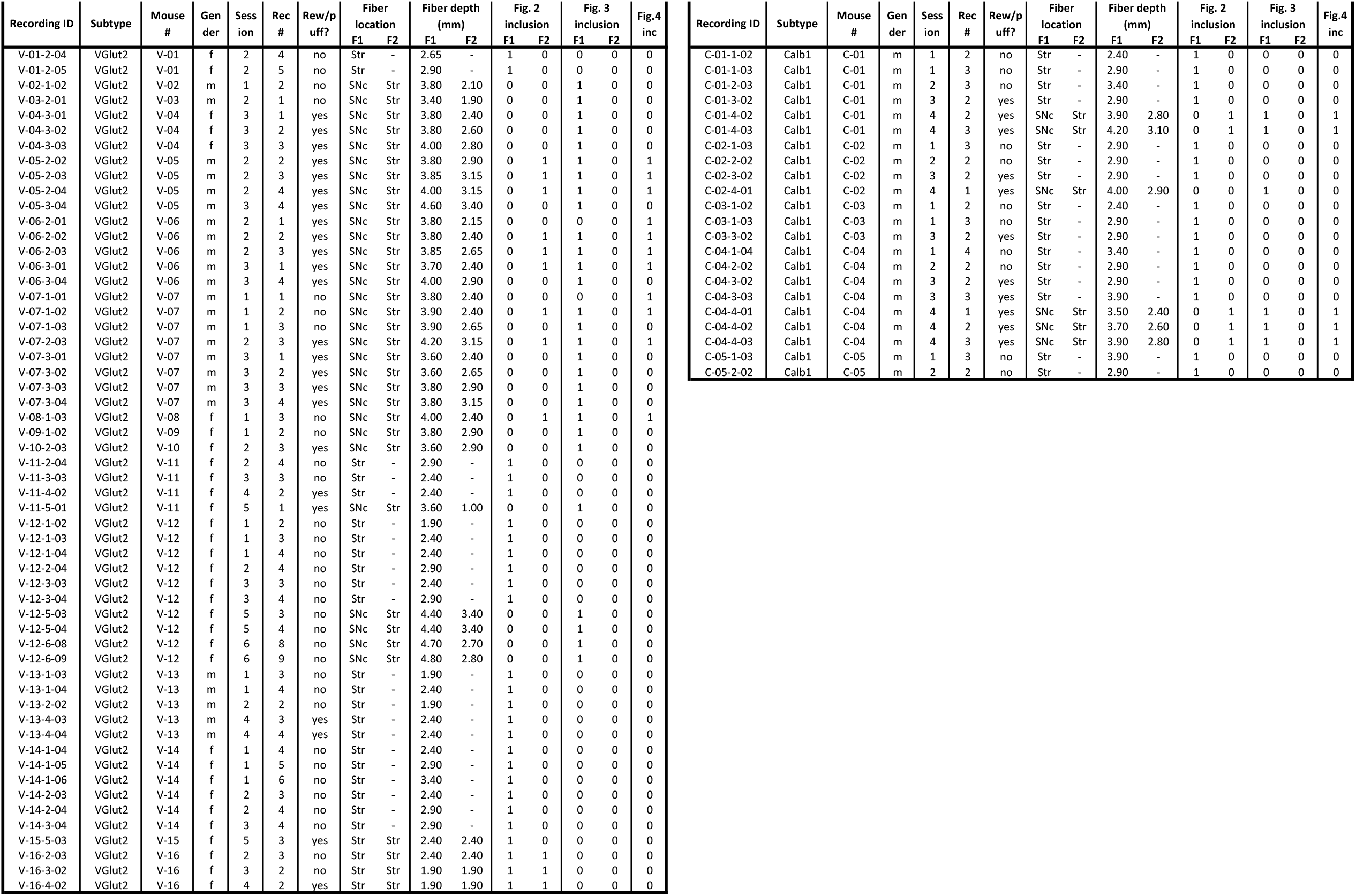
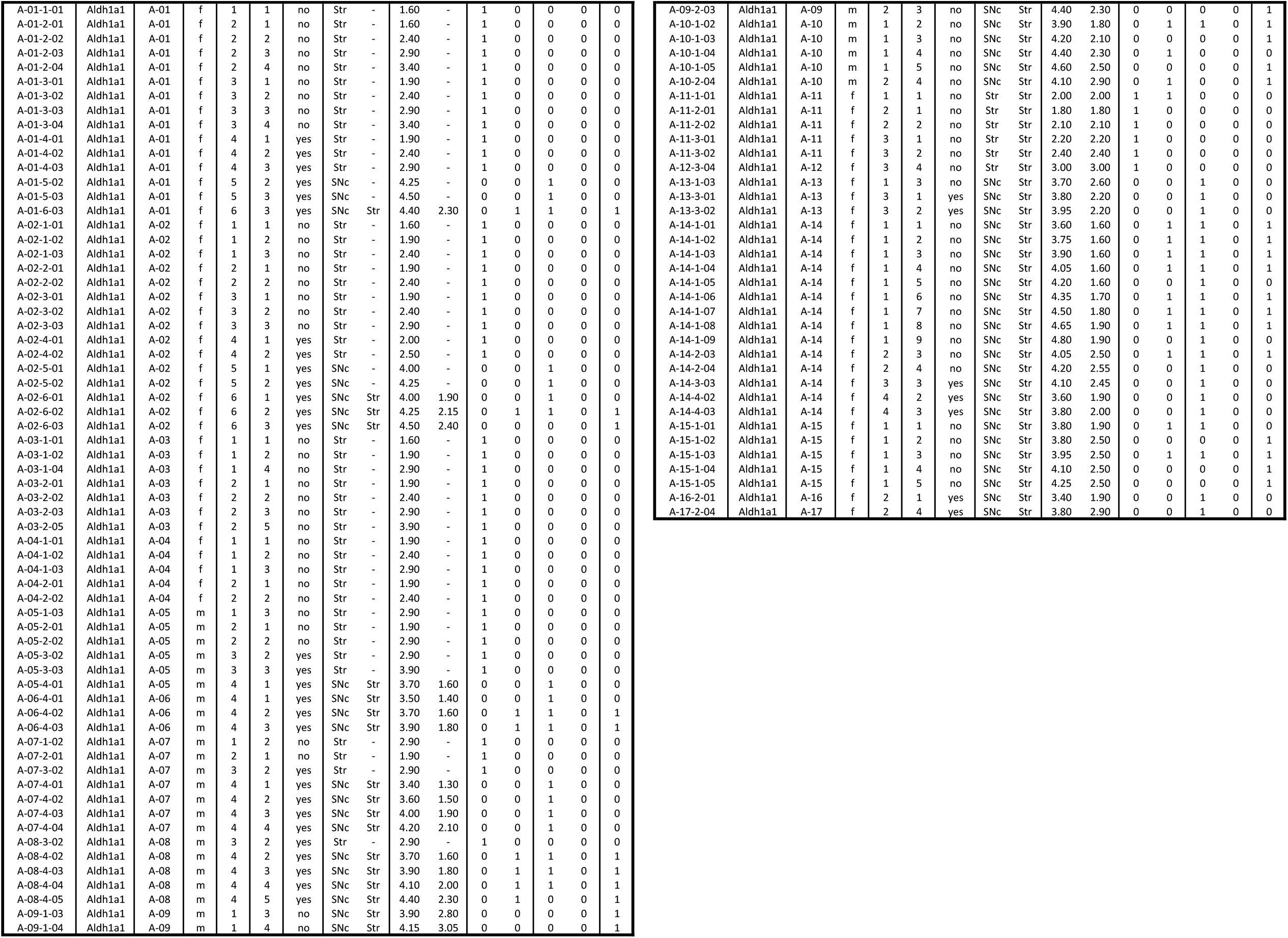
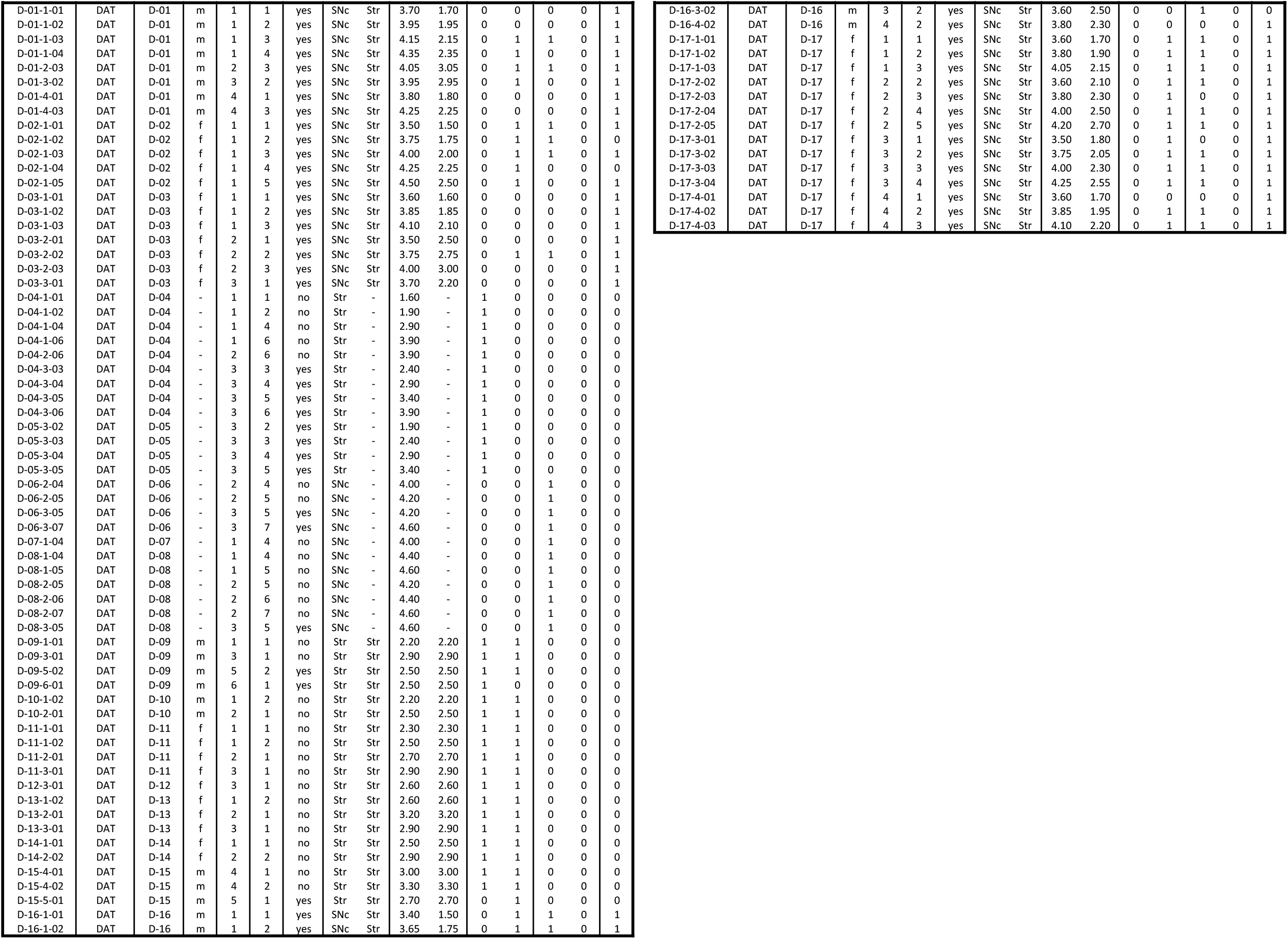
Metadata for each recording included in this paper divided by subtype. One or two fibers (F1,F2) were used for each recording. Rightmost columns show what figures each recording was included in.

| Recording ID | Subtype | Mouse # | Gender | Session | Rec # | Rew/puff? | Fiber location |  | Fiber depth (mm) |  | Fig. 2 inclusion |  | Fig. 3 inclusion |  | Fig.4 inc |
| --- | --- | --- | --- | --- | --- | --- | --- | --- | --- | --- | --- | --- | --- | --- | --- |
|  |  |  |  |  |  |  | F1 | F2 | F1 | F2 | F1 | F2 | F1 | F2 |  |
| V-01-2-04 | VGlut2 | V-01 | f | 2 | 4 | no | Str | - | 2.65 | - | 1 | 0 | 0 | 0 | 0 |
| V-01-2-05 | VGlut2 | V-01 | f | 2 | 5 | no | Str | - | 2.90 | - | 1 | 0 | 0 | 0 | 0 |
| V-02-1-02 | VGlut2 | V-02 | m | 1 | 2 | no | SNc | Str | 3.80 | 2.10 | 0 | 0 | 1 | 0 | 0 |
| V-03-2-01 | VGlut2 | V-03 | m | 2 | 1 | no | SNc | Str | 3.40 | 1.90 | 0 | 0 | 1 | 0 | 0 |
| V-04-3-01 | VGlut2 | V-04 | f | 3 | 1 | yes | SNc | Str | 3.80 | 2.40 | 0 | 0 | 1 | 0 | 0 |
| V-04-3-02 | VGlut2 | V-04 | f | 3 | 2 | yes | SNc | Str | 3.80 | 2.60 | 0 | 0 | 1 | 0 | 0 |
| V-04-3-03 | VGlut2 | V-04 | f | 3 | 3 | yes | SNc | Str | 4.00 | 2.80 | 0 | 0 | 1 | 0 | 0 |
| V-05-2-02 | VGlut2 | V-05 | m | 2 | 2 | yes | SNc | Str | 3.80 | 2.90 | 0 | 1 | 1 | 0 | 1 |
| V-05-2-03 | VGlut2 | V-05 | m | 2 | 3 | yes | SNc | Str | 3.85 | 3.15 | 0 | 1 | 1 | 0 | 1 |
| V-05-2-04 | VGlut2 | V-05 | m | 2 | 4 | yes | SNc | Str | 4.00 | 3.15 | 0 | 1 | 1 | 0 | 1 |
| V-05-3-04 | VGlut2 | V-05 | m | 3 | 4 | yes | SNc | Str | 4.60 | 3.40 | 0 | 0 | 1 | 0 | 0 |
| V-06-2-01 | VGlut2 | V-06 | m | 2 | 1 | yes | SNc | Str | 3.80 | 2.15 | 0 | 0 | 0 | 0 | 1 |
| V-06-2-02 | VGlut2 | V-06 | m | 2 | 2 | yes | SNc | Str | 3.80 | 2.40 | 0 | 1 | 1 | 0 | 1 |
| V-06-2-03 | VGlut2 | V-06 | m | 2 | 3 | yes | SNc | Str | 3.85 | 2.65 | 0 | 1 | 1 | 0 | 1 |
| V-06-3-01 | VGlut2 | V-06 | m | 3 | 1 | yes | SNc | Str | 3.70 | 2.40 | 0 | 1 | 1 | 0 | 1 |
| V-06-3-04 | VGlut2 | V-06 | m | 3 | 4 | yes | SNc | Str | 4.00 | 2.90 | 0 | 0 | 1 | 0 | 0 |
| V-07-1-01 | VGlut2 | V-07 | m | 1 | 1 | no | SNc | Str | 3.80 | 2.40 | 0 | 0 | 0 | 0 | 1 |
| V-07-1-02 | VGlut2 | V-07 | m | 1 | 2 | no | SNc | Str | 3.90 | 2.40 | 0 | 1 | 1 | 0 | 1 |
| V-07-1-03 | VGlut2 | V-07 | m | 1 | 3 | no | SNc | Str | 3.90 | 2.65 | 0 | 0 | 1 | 0 | 0 |
| V-07-2-03 | VGlut2 | V-07 | m | 2 | 3 | yes | SNc | Str | 4.20 | 3.15 | 0 | 1 | 1 | 0 | 1 |
| V-07-3-01 | VGlut2 | V-07 | m | 3 | 1 | yes | SNc | Str | 3.60 | 2.40 | 0 | 0 | 1 | 0 | 0 |
| V-07-3-02 | VGlut2 | V-07 | m | 3 | 2 | yes | SNc | Str | 3.60 | 2.65 | 0 | 0 | 1 | 0 | 0 |
| V-07-3-03 | VGlut2 | V-07 | m | 3 | 3 | yes | SNc | Str | 3.80 | 2.90 | 0 | 0 | 1 | 0 | 0 |
| V-07-3-04 | VGlut2 | V-07 | m | 3 | 4 | yes | SNc | Str | 3.80 | 3.15 | 0 | 0 | 1 | 0 | 0 |
| V-08-1-03 | VGlut2 | V-08 | f | 1 | 3 | no | SNc | Str | 4.00 | 2.40 | 0 | 1 | 1 | 0 | 1 |
| V-09-1-02 | VGlut2 | V-09 | f | 1 | 2 | no | SNc | Str | 3.80 | 2.90 | 0 | 0 | 1 | 0 | 0 |
| V-10-2-03 | VGlut2 | V-10 | f | 2 | 3 | yes | SNc | Str | 3.60 | 2.90 | 0 | 0 | 1 | 0 | 0 |
| V-11-2-04 | VGlut2 | V-11 | f | 2 | 4 | no | Str | - | 2.90 | - | 1 | 0 | 0 | 0 | 0 |
| V-11-3-03 | VGlut2 | V-11 | f | 3 | 3 | no | Str | - | 2.40 | - | 1 | 0 | 0 | 0 | 0 |
| V-11-4-02 | VGlut2 | V-11 | f | 4 | 2 | yes | Str | - | 2.40 | - | 1 | 0 | 0 | 0 | 0 |
| V-11-5-01 | VGlut2 | V-11 | f | 5 | 1 | yes | SNc | Str | 3.60 | 1.00 | 0 | 0 | 1 | 0 | 0 |
| V-12-1-02 | VGlut2 | V-12 | f | 1 | 2 | no | Str | - | 1.90 | - | 1 | 0 | 0 | 0 | 0 |
| V-12-1-03 | VGlut2 | V-12 | f | 1 | 3 | no | Str | - | 2.40 | - | 1 | 0 | 0 | 0 | 0 |
| V-12-1-04 | VGlut2 | V-12 | f | 1 | 4 | no | Str | - | 2.40 | - | 1 | 0 | 0 | 0 | 0 |
| V-12-2-04 | VGlut2 | V-12 | f | 2 | 4 | no | Str | - | 2.90 | - | 1 | 0 | 0 | 0 | 0 |
| V-12-3-03 | VGlut2 | V-12 | f | 3 | 3 | no | Str | - | 2.40 | - | 1 | 0 | 0 | 0 | 0 |
| V-12-3-04 | VGlut2 | V-12 | f | 3 | 4 | no | Str | - | 2.90 | - | 1 | 0 | 0 | 0 | 0 |
| V-12-5-03 | VGlut2 | V-12 | f | 5 | 3 | no | SNc | Str | 4.40 | 3.40 | 0 | 0 | 1 | 0 | 0 |
| V-12-5-04 | VGlut2 | V-12 | f | 5 | 4 | no | SNc | Str | 4.40 | 3.40 | 0 | 0 | 1 | 0 | 0 |
| V-12-6-08 | VGlut2 | V-12 | f | 6 | 8 | no | SNc | Str | 4.70 | 2.70 | 0 | 0 | 1 | 0 | 0 |
| V-12-6-09 | VGlut2 | V-12 | f | 6 | 9 | no | SNc | Str | 4.80 | 2.80 | 0 | 0 | 1 | 0 | 0 |
| V-13-1-03 | VGlut2 | V-13 | m | 1 | 3 | no | Str | - | 1.90 | - | 1 | 0 | 0 | 0 | 0 |
| V-13-1-04 | VGlut2 | V-13 | m | 1 | 4 | no | Str | - | 2.40 | - | 1 | 0 | 0 | 0 | 0 |
| V-13-2-02 | VGlut2 | V-13 | m | 2 | 2 | no | Str | - | 1.90 | - | 1 | 0 | 0 | 0 | 0 |
| V-13-4-03 | VGlut2 | V-13 | m | 4 | 3 | yes | Str | - | 2.40 | - | 1 | 0 | 0 | 0 | 0 |
| V-13-4-04 | VGlut2 | V-13 | m | 4 | 4 | yes | Str | - | 2.40 | - | 1 | 0 | 0 | 0 | 0 |
| V-14-1-04 | VGlut2 | V-14 | f | 1 | 4 | no | Str | - | 2.40 | - | 1 | 0 | 0 | 0 | 0 |
| V-14-1-05 | VGlut2 | V-14 | f | 1 | 5 | no | Str | - | 2.90 | - | 1 | 0 | 0 | 0 | 0 |
| V-14-1-06 | VGlut2 | V-14 | f | 1 | 6 | no | Str | - | 3.40 | - | 1 | 0 | 0 | 0 | 0 |
| V-14-2-03 | VGlut2 | V-14 | f | 2 | 3 | no | Str | - | 2.40 | - | 1 | 0 | 0 | 0 | 0 |
| V-14-2-04 | VGlut2 | V-14 | f | 2 | 4 | no | Str | - | 2.90 | - | 1 | 0 | 0 | 0 | 0 |
| V-14-3-04 | VGlut2 | V-14 | f | 3 | 4 | no | Str | - | 2.90 | - | 1 | 0 | 0 | 0 | 0 |
| V-15-5-03 | VGlut2 | V-15 | f | 5 | 3 | yes | Str | Str | 2.40 | 2.40 | 1 | 0 | 0 | 0 | 0 |
| V-16-2-03 | VGlut2 | V-16 | f | 2 | 3 | no | Str | Str | 2.40 | 2.40 | 1 | 1 | 0 | 0 | 0 |
| V-16-3-02 | VGlut2 | V-16 | f | 3 | 2 | no | Str | Str | 1.90 | 1.90 | 1 | 1 | 0 | 0 | 0 |
| V-16-4-02 | VGlut2 | V-16 | f | 4 | 2 | yes | Str | Str | 1.90 | 1.90 | 1 | 1 | 0 | 0 | 0 |
| C-01-1-02 | Calb1 | C-01 | m | 1 | 2 | no | Str | - | 2.40 | - | 1 | 0 | 0 | 0 | 0 |
| C-01-1-03 | Calb1 | C-01 | m | 1 | 3 | no | Str | - | 2.90 | - | 1 | 0 | 0 | 0 | 0 |
| C-01-2-03 | Calb1 | C-01 | m | 2 | 3 | no | Str | - | 3.40 | - | 1 | 0 | 0 | 0 | 0 |
| C-01-3-02 | Calb1 | C-01 | m | 3 | 2 | yes | Str | - | 2.90 | - | 1 | 0 | 0 | 0 | 0 |
| C-01-4-02 | Calb1 | C-01 | m | 4 | 2 | yes | SNc | Str | 3.90 | 2.80 | 0 | 1 | 1 | 0 | 1 |
| C-01-4-03 | Calb1 | C-01 | m | 4 | 3 | yes | SNc | Str | 4.20 | 3.10 | 0 | 1 | 1 | 0 | 1 |
| C-02-1-03 | Calb1 | C-02 | m | 1 | 3 | no | Str | - | 2.90 | - | 1 | 0 | 0 | 0 | 0 |
| C-02-2-02 | Calb1 | C-02 | m | 2 | 2 | no | Str | - | 2.90 | - | 1 | 0 | 0 | 0 | 0 |
| C-02-3-02 | Calb1 | C-02 | m | 3 | 2 | yes | Str | - | 2.90 | - | 1 | 0 | 0 | 0 | 0 |
| C-02-4-01 | Calb1 | C-02 | m | 4 | 1 | yes | SNc | Str | 4.00 | 2.90 | 0 | 0 | 1 | 0 | 0 |
| C-03-1-02 | Calb1 | C-03 | m | 1 | 2 | no | Str | - | 2.40 | - | 1 | 0 | 0 | 0 | 0 |
| C-03-1-03 | Calb1 | C-03 | m | 1 | 3 | no | Str | - | 2.90 | - | 1 | 0 | 0 | 0 | 0 |
| C-03-3-02 | Calb1 | C-03 | m | 3 | 2 | yes | Str | - | 2.90 | - | 1 | 0 | 0 | 0 | 0 |
| C-04-1-04 | Calb1 | C-04 | m | 1 | 4 | no | Str | - | 3.40 | - | 1 | 0 | 0 | 0 | 0 |
| C-04-2-02 | Calb1 | C-04 | m | 2 | 2 | no | Str | - | 2.90 | - | 1 | 0 | 0 | 0 | 0 |
| C-04-3-02 | Calb1 | C-04 | m | 3 | 2 | yes | Str | - | 2.90 | - | 1 | 0 | 0 | 0 | 0 |
| C-04-3-03 | Calb1 | C-04 | m | 3 | 3 | yes | Str | - | 3.90 | - | 1 | 0 | 0 | 0 | 0 |
| C-04-4-01 | Calb1 | C-04 | m | 4 | 1 | yes | SNc | Str | 3.50 | 2.40 | 0 | 1 | 1 | 0 | 1 |
| C-04-4-02 | Calb1 | C-04 | m | 4 | 2 | yes | SNc | Str | 3.70 | 2.60 | 0 | 1 | 1 | 0 | 1 |
| C-04-4-03 | Calb1 | C-04 | m | 4 | 3 | yes | SNc | Str | 3.90 | 2.80 | 0 | 1 | 1 | 0 | 1 |
| C-05-1-03 | Calb1 | C-05 | m | 1 | 3 | no | Str | - | 3.90 | - | 1 | 0 | 0 | 0 | 0 |
| C-05-2-02 | Calb1 | C-05 | m | 2 | 2 | no | Str | - | 2.90 | - | 1 | 0 | 0 | 0 | 0 |

| Recording ID | Subtype | Mouse # | Gen der | Sess ion | Rec # | Rew/p uff? | Fiber location<br>F1 F2 | Fiber depth<br>(mm)<br>F1 F2 | Fig. 2<br>inclusion<br>F1 F2 | Fig. 3<br>inclusion<br>F1 F2 | Fig.4<br>inc |
| --- | --- | --- | --- | --- | --- | --- | --- | --- | --- | --- | --- |
| A-01-1-01 | Aldh1a1 | A-01 | f | 1 | 1 | no | Str - | 1.60 - | 1 0 | 0 0 | 0 |
| A-01-2-01 | Aldh1a1 | A-01 | f | 2 | 1 | no | Str - | 1.60 - | 1 0 | 0 0 | 0 |
| A-01-2-02 | Aldh1a1 | A-01 | f | 2 | 2 | no | Str - | 2.40 - | 1 0 | 0 0 | 0 |
| A-01-2-03 | Aldh1a1 | A-01 | f | 2 | 3 | no | Str - | 2.90 - | 1 0 | 0 0 | 0 |
| A-01-2-04 | Aldh1a1 | A-01 | f | 2 | 4 | no | Str - | 3.40 - | 1 0 | 0 0 | 0 |
| A-01-3-01 | Aldh1a1 | A-01 | f | 3 | 1 | no | Str - | 1.90 - | 1 0 | 0 0 | 0 |
| A-01-3-02 | Aldh1a1 | A-01 | f | 3 | 2 | no | Str - | 2.40 - | 1 0 | 0 0 | 0 |
| A-01-3-03 | Aldh1a1 | A-01 | f | 3 | 3 | no | Str - | 2.90 - | 1 0 | 0 0 | 0 |
| A-01-3-04 | Aldh1a1 | A-01 | f | 3 | 4 | no | Str - | 3.40 - | 1 0 | 0 0 | 0 |
| A-01-4-01 | Aldh1a1 | A-01 | f | 4 | 1 | yes | Str - | 1.90 - | 1 0 | 0 0 | 0 |
| A-01-4-02 | Aldh1a1 | A-01 | f | 4 | 2 | yes | Str - | 2.40 - | 1 0 | 0 0 | 0 |
| A-01-4-03 | Aldh1a1 | A-01 | f | 4 | 3 | yes | Str - | 2.90 - | 1 0 | 0 0 | 0 |
| A-01-5-02 | Aldh1a1 | A-01 | f | 5 | 2 | yes | SNc - | 4.25 - | 0 0 | 1 0 | 0 |
| A-01-5-03 | Aldh1a1 | A-01 | f | 5 | 3 | yes | SNc - | 4.50 - | 0 0 | 1 0 | 0 |
| A-01-6-03 | Aldh1a1 | A-01 | f | 6 | 3 | yes | SNc Str | 4.40 2.30 | 0 1 | 1 0 | 1 |
| A-02-1-01 | Aldh1a1 | A-02 | f | 1 | 1 | no | Str - | 1.60 - | 1 0 | 0 0 | 0 |
| A-02-1-02 | Aldh1a1 | A-02 | f | 1 | 2 | no | Str - | 1.90 - | 1 0 | 0 0 | 0 |
| A-02-1-03 | Aldh1a1 | A-02 | f | 1 | 3 | no | Str - | 2.40 - | 1 0 | 0 0 | 0 |
| A-02-2-01 | Aldh1a1 | A-02 | f | 2 | 1 | no | Str - | 1.90 - | 1 0 | 0 0 | 0 |
| A-02-2-02 | Aldh1a1 | A-02 | f | 2 | 2 | no | Str - | 2.40 - | 1 0 | 0 0 | 0 |
| A-02-3-01 | Aldh1a1 | A-02 | f | 3 | 1 | no | Str - | 1.90 - | 1 0 | 0 0 | 0 |
| A-02-3-02 | Aldh1a1 | A-02 | f | 3 | 2 | no | Str - | 2.40 - | 1 0 | 0 0 | 0 |
| A-02-3-03 | Aldh1a1 | A-02 | f | 3 | 3 | no | Str - | 2.90 - | 1 0 | 0 0 | 0 |
| A-02-4-01 | Aldh1a1 | A-02 | f | 4 | 1 | yes | Str - | 2.00 - | 1 0 | 0 0 | 0 |
| A-02-4-02 | Aldh1a1 | A-02 | f | 4 | 2 | yes | Str - | 2.50 - | 1 0 | 0 0 | 0 |
| A-02-5-01 | Aldh1a1 | A-02 | f | 5 | 1 | yes | SNc - | 4.00 - | 0 0 | 1 0 | 0 |
| A-02-5-02 | Aldh1a1 | A-02 | f | 5 | 2 | yes | SNc - | 4.25 - | 0 0 | 1 0 | 0 |
| A-02-6-01 | Aldh1a1 | A-02 | f | 6 | 1 | yes | SNc Str | 4.00 1.90 | 0 0 | 1 0 | 0 |
| A-02-6-02 | Aldh1a1 | A-02 | f | 6 | 2 | yes | SNc Str | 4.25 2.15 | 0 1 | 1 0 | 1 |
| A-02-6-03 | Aldh1a1 | A-02 | f | 6 | 3 | yes | SNc Str | 4.50 2.40 | 0 0 | 0 0 | 1 |
| A-03-1-01 | Aldh1a1 | A-03 | f | 1 | 1 | no | Str - | 1.60 - | 1 0 | 0 0 | 0 |
| A-03-1-02 | Aldh1a1 | A-03 | f | 1 | 2 | no | Str - | 1.90 - | 1 0 | 0 0 | 0 |
| A-03-1-04 | Aldh1a1 | A-03 | f | 1 | 4 | no | Str - | 2.90 - | 1 0 | 0 0 | 0 |
| A-03-2-01 | Aldh1a1 | A-03 | f | 2 | 1 | no | Str - | 1.90 - | 1 0 | 0 0 | 0 |
| A-03-2-02 | Aldh1a1 | A-03 | f | 2 | 2 | no | Str - | 2.40 - | 1 0 | 0 0 | 0 |
| A-03-2-03 | Aldh1a1 | A-03 | f | 2 | 3 | no | Str - | 2.90 - | 1 0 | 0 0 | 0 |
| A-03-2-05 | Aldh1a1 | A-03 | f | 2 | 5 | no | Str - | 3.90 - | 1 0 | 0 0 | 0 |
| A-04-1-01 | Aldh1a1 | A-04 | f | 1 | 1 | no | Str - | 1.90 - | 1 0 | 0 0 | 0 |
| A-04-1-02 | Aldh1a1 | A-04 | f | 1 | 2 | no | Str - | 2.40 - | 1 0 | 0 0 | 0 |
| A-04-1-03 | Aldh1a1 | A-04 | f | 1 | 3 | no | Str - | 2.90 - | 1 0 | 0 0 | 0 |
| A-04-2-01 | Aldh1a1 | A-04 | f | 2 | 1 | no | Str - | 1.90 - | 1 0 | 0 0 | 0 |
| A-04-2-02 | Aldh1a1 | A-04 | f | 2 | 2 | no | Str - | 2.40 - | 1 0 | 0 0 | 0 |
| A-05-1-03 | Aldh1a1 | A-05 | m | 1 | 3 | no | Str - | 2.90 - | 1 0 | 0 0 | 0 |
| A-05-2-01 | Aldh1a1 | A-05 | m | 2 | 1 | no | Str - | 1.90 - | 1 0 | 0 0 | 0 |
| A-05-2-02 | Aldh1a1 | A-05 | m | 2 | 2 | no | Str - | 2.90 - | 1 0 | 0 0 | 0 |
| A-05-3-02 | Aldh1a1 | A-05 | m | 3 | 2 | yes | Str - | 2.90 - | 1 0 | 0 0 | 0 |
| A-05-3-03 | Aldh1a1 | A-05 | m | 3 | 3 | yes | Str - | 3.90 - | 1 0 | 0 0 | 0 |
| A-05-4-01 | Aldh1a1 | A-05 | m | 4 | 1 | yes | SNc Str | 3.70 1.60 | 0 0 | 1 0 | 0 |
| A-06-4-01 | Aldh1a1 | A-06 | m | 4 | 1 | yes | SNc Str | 3.50 1.40 | 0 0 | 1 0 | 0 |
| A-06-4-02 | Aldh1a1 | A-06 | m | 4 | 2 | yes | SNc Str | 3.70 1.60 | 0 1 | 1 0 | 1 |
| A-06-4-03 | Aldh1a1 | A-06 | m | 4 | 3 | yes | SNc Str | 3.90 1.80 | 0 1 | 1 0 | 1 |
| A-07-1-02 | Aldh1a1 | A-07 | m | 1 | 2 | no | Str - | 2.90 - | 1 0 | 0 0 | 0 |
| A-07-2-01 | Aldh1a1 | A-07 | m | 2 | 1 | no | Str - | 1.90 - | 1 0 | 0 0 | 0 |
| A-07-3-02 | Aldh1a1 | A-07 | m | 3 | 2 | yes | Str - | 2.90 - | 1 0 | 0 0 | 0 |
| A-07-4-01 | Aldh1a1 | A-07 | m | 4 | 1 | yes | SNc Str | 3.40 1.30 | 0 0 | 1 0 | 0 |
| A-07-4-02 | Aldh1a1 | A-07 | m | 4 | 2 | yes | SNc Str | 3.60 1.50 | 0 0 | 1 0 | 0 |
| A-07-4-03 | Aldh1a1 | A-07 | m | 4 | 3 | yes | SNc Str | 4.00 1.90 | 0 0 | 1 0 | 0 |
| A-07-4-04 | Aldh1a1 | A-07 | m | 4 | 4 | yes | SNc Str | 4.20 2.10 | 0 0 | 1 0 | 0 |
| A-08-3-02 | Aldh1a1 | A-08 | m | 3 | 2 | yes | Str - | 2.90 - | 1 0 | 0 0 | 0 |
| A-08-4-02 | Aldh1a1 | A-08 | m | 4 | 2 | yes | SNc Str | 3.70 1.60 | 0 1 | 1 0 | 1 |
| A-08-4-03 | Aldh1a1 | A-08 | m | 4 | 3 | yes | SNc Str | 3.90 1.80 | 0 1 | 1 0 | 1 |
| A-08-4-04 | Aldh1a1 | A-08 | m | 4 | 4 | yes | SNc Str | 4.10 2.00 | 0 1 | 1 0 | 1 |
| A-08-4-05 | Aldh1a1 | A-08 | m | 4 | 5 | yes | SNc Str | 4.40 2.30 | 0 1 | 0 0 | 1 |
| A-09-1-03 | Aldh1a1 | A-09 | m | 1 | 3 | no | SNc Str | 3.90 2.80 | 0 0 | 0 0 | 1 |
| A-09-1-04 | Aldh1a1 | A-09 | m | 1 | 4 | no | SNc Str | 4.15 3.05 | 0 0 | 0 0 | 1 |
| A-09-2-03 | Aldh1a1 | A-09 | m | 2 | 3 | no | SNc Str | 4.40 2.30 | 0 0 | 0 0 | 1 |
| A-10-1-02 | Aldh1a1 | A-10 | m | 1 | 2 | no | SNc Str | 3.90 1.80 | 0 1 | 1 0 | 1 |
| A-10-1-03 | Aldh1a1 | A-10 | m | 1 | 3 | no | SNc Str | 4.20 2.10 | 0 0 | 0 0 | 1 |
| A-10-1-04 | Aldh1a1 | A-10 | m | 1 | 4 | no | SNc Str | 4.40 2.30 | 0 1 | 0 0 | 0 |
| A-10-1-05 | Aldh1a1 | A-10 | m | 1 | 5 | no | SNc Str | 4.60 2.50 | 0 0 | 0 0 | 1 |
| A-10-2-04 | Aldh1a1 | A-10 | m | 2 | 4 | no | SNc Str | 4.10 2.90 | 0 1 | 0 0 | 1 |
| A-11-1-01 | Aldh1a1 | A-11 | f | 1 | 1 | no | Str Str | 2.00 2.00 | 1 1 | 0 0 | 0 |
| A-11-2-01 | Aldh1a1 | A-11 | f | 2 | 1 | no | Str Str | 1.80 1.80 | 1 0 | 0 0 | 0 |
| A-11-2-02 | Aldh1a1 | A-11 | f | 2 | 2 | no | Str Str | 2.10 2.10 | 1 0 | 0 0 | 0 |
| A-11-3-01 | Aldh1a1 | A-11 | f | 3 | 1 | no | Str Str | 2.20 2.20 | 1 0 | 0 0 | 0 |
| A-11-3-02 | Aldh1a1 | A-11 | f | 3 | 2 | no | Str Str | 2.40 2.40 | 1 0 | 0 0 | 0 |
| A-12-3-04 | Aldh1a1 | A-12 | f | 3 | 4 | no | Str Str | 3.00 3.00 | 1 0 | 0 0 | 0 |
| A-13-1-03 | Aldh1a1 | A-13 | f | 1 | 3 | no | SNc Str | 3.70 2.60 | 0 0 | 1 0 | 0 |
| A-13-3-01 | Aldh1a1 | A-13 | f | 3 | 1 | yes | SNc Str | 3.80 2.20 | 0 0 | 1 0 | 0 |
| A-13-3-02 | Aldh1a1 | A-13 | f | 3 | 2 | yes | SNc Str | 3.95 2.20 | 0 0 | 1 0 | 0 |
| A-14-1-01 | Aldh1a1 | A-14 | f | 1 | 1 | no | SNc Str | 3.60 1.60 | 0 1 | 1 0 | 1 |
| A-14-1-02 | Aldh1a1 | A-14 | f | 1 | 2 | no | SNc Str | 3.75 1.60 | 0 1 | 1 0 | 1 |
| A-14-1-03 | Aldh1a1 | A-14 | f | 1 | 3 | no | SNc Str | 3.90 1.60 | 0 1 | 1 0 | 1 |
| A-14-1-04 | Aldh1a1 | A-14 | f | 1 | 4 | no | SNc Str | 4.05 1.60 | 0 1 | 1 0 | 1 |
| A-14-1-05 | Aldh1a1 | A-14 | f | 1 | 5 | no | SNc Str | 4.20 1.60 | 0 0 | 1 0 | 0 |
| A-14-1-06 | Aldh1a1 | A-14 | f | 1 | 6 | no | SNc Str | 4.35 1.70 | 0 1 | 1 0 | 1 |
| A-14-1-07 | Aldh1a1 | A-14 | f | 1 | 7 | no | SNc Str | 4.50 1.80 | 0 1 | 1 0 | 1 |
| A-14-1-08 | Aldh1a1 | A-14 | f | 1 | 8 | no | SNc Str | 4.65 1.90 | 0 1 | 1 0 | 1 |
| A-14-1-09 | Aldh1a1 | A-14 | f | 1 | 9 | no | SNc Str | 4.80 1.90 | 0 0 | 1 0 | 0 |
| A-14-2-03 | Aldh1a1 | A-14 | f | 2 | 3 | no | SNc Str | 4.05 2.50 | 0 1 | 1 0 | 1 |
| A-14-2-04 | Aldh1a1 | A-14 | f | 2 | 4 | no | SNc Str | 4.20 2.55 | 0 0 | 1 0 | 0 |
| A-14-3-03 | Aldh1a1 | A-14 | f | 3 | 3 | yes | SNc Str | 4.10 2.45 | 0 0 | 1 0 | 0 |
| A-14-4-02 | Aldh1a1 | A-14 | f | 4 | 2 | yes | SNc Str | 3.60 1.90 | 0 0 | 1 0 | 0 |
| A-14-4-03 | Aldh1a1 | A-14 | f | 4 | 3 | yes | SNc Str | 3.80 2.00 | 0 0 | 1 0 | 0 |
| A-15-1-01 | Aldh1a1 | A-15 | f | 1 | 1 | no | SNc Str | 3.80 1.90 | 0 1 | 1 0 | 0 |
| A-15-1-02 | Aldh1a1 | A-15 | f | 1 | 2 | no | SNc Str | 3.80 2.50 | 0 0 | 0 0 | 1 |
| A-15-1-03 | Aldh1a1 | A-15 | f | 1 | 3 | no | SNc Str | 3.95 2.50 | 0 1 | 1 0 | 1 |
| A-15-1-04 | Aldh1a1 | A-15 | f | 1 | 4 | no | SNc Str | 4.10 2.50 | 0 0 | 0 0 | 1 |
| A-15-1-05 | Aldh1a1 | A-15 | f | 1 | 5 | no | SNc Str | 4.25 2.50 | 0 0 | 0 0 | 1 |
| A-16-2-01 | Aldh1a1 | A-16 | f | 2 | 1 | yes | SNc Str | 3.40 1.90 | 0 0 | 1 0 | 0 |
| A-17-2-04 | Aldh1a1 | A-17 | f | 2 | 4 | yes | SNc Str | 3.80 2.90 | 0 0 | 1 0 | 0 |
|  |  |  |  |  |  |  | F1 | F2 | F1 | F2 | F1 | F2 | F1 | F2 |  |
| D-01-1-01 | DAT | D-01 | m | 1 | 1 | yes | SNc | Str | 3.70 | 1.70 | 0 | 0 | 0 | 0 | 1 |
| D-01-1-02 | DAT | D-01 | m | 1 | 2 | yes | SNc | Str | 3.95 | 1.95 | 0 | 0 | 0 | 0 | 1 |
| D-01-1-03 | DAT | D-01 | m | 1 | 3 | yes | SNc | Str | 4.15 | 2.15 | 0 | 1 | 1 | 0 | 1 |
| D-01-1-04 | DAT | D-01 | m | 1 | 4 | yes | SNc | Str | 4.35 | 2.35 | 0 | 1 | 0 | 0 | 1 |
| D-01-2-03 | DAT | D-01 | m | 2 | 3 | yes | SNc | Str | 4.05 | 3.05 | 0 | 1 | 1 | 0 | 1 |
| D-01-3-02 | DAT | D-01 | m | 3 | 2 | yes | SNc | Str | 3.95 | 2.95 | 0 | 1 | 0 | 0 | 1 |
| D-01-4-01 | DAT | D-01 | m | 4 | 1 | yes | SNc | Str | 3.80 | 1.80 | 0 | 0 | 0 | 0 | 1 |
| D-01-4-03 | DAT | D-01 | m | 4 | 3 | yes | SNc | Str | 4.25 | 2.25 | 0 | 0 | 0 | 0 | 1 |
| D-02-1-01 | DAT | D-02 | f | 1 | 1 | yes | SNc | Str | 3.50 | 1.50 | 0 | 1 | 1 | 0 | 1 |
| D-02-1-02 | DAT | D-02 | f | 1 | 2 | yes | SNc | Str | 3.75 | 1.75 | 0 | 1 | 1 | 0 | 0 |
| D-02-1-03 | DAT | D-02 | f | 1 | 3 | yes | SNc | Str | 4.00 | 2.00 | 0 | 1 | 1 | 0 | 1 |
| D-02-1-04 | DAT | D-02 | f | 1 | 4 | yes | SNc | Str | 4.25 | 2.25 | 0 | 1 | 0 | 0 | 0 |
| D-02-1-05 | DAT | D-02 | f | 1 | 5 | yes | SNc | Str | 4.50 | 2.50 | 0 | 1 | 0 | 0 | 1 |
| D-03-1-01 | DAT | D-03 | f | 1 | 1 | yes | SNc | Str | 3.60 | 1.60 | 0 | 0 | 0 | 0 | 1 |
| D-03-1-02 | DAT | D-03 | f | 1 | 2 | yes | SNc | Str | 3.85 | 1.85 | 0 | 0 | 0 | 0 | 1 |
| D-03-1-03 | DAT | D-03 | f | 1 | 3 | yes | SNc | Str | 4.10 | 2.10 | 0 | 0 | 0 | 0 | 1 |
| D-03-2-01 | DAT | D-03 | f | 2 | 1 | yes | SNc | Str | 3.50 | 2.50 | 0 | 0 | 0 | 0 | 1 |
| D-03-2-02 | DAT | D-03 | f | 2 | 2 | yes | SNc | Str | 3.75 | 2.75 | 0 | 1 | 1 | 0 | 1 |
| D-03-2-03 | DAT | D-03 | f | 2 | 3 | yes | SNc | Str | 4.00 | 3.00 | 0 | 0 | 0 | 0 | 1 |
| D-03-3-01 | DAT | D-03 | f | 3 | 1 | yes | SNc | Str | 3.70 | 2.20 | 0 | 0 | 0 | 0 | 1 |
| D-04-1-01 | DAT | D-04 | - | 1 | 1 | no | Str | - | 1.60 | - | 1 | 0 | 0 | 0 | 0 |
| D-04-1-02 | DAT | D-04 | - | 1 | 2 | no | Str | - | 1.90 | - | 1 | 0 | 0 | 0 | 0 |
| D-04-1-04 | DAT | D-04 | - | 1 | 4 | no | Str | - | 2.90 | - | 1 | 0 | 0 | 0 | 0 |
| D-04-1-06 | DAT | D-04 | - | 1 | 6 | no | Str | - | 3.90 | - | 1 | 0 | 0 | 0 | 0 |
| D-04-2-06 | DAT | D-04 | - | 2 | 6 | no | Str | - | 3.90 | - | 1 | 0 | 0 | 0 | 0 |
| D-04-3-03 | DAT | D-04 | - | 3 | 3 | yes | Str | - | 2.40 | - | 1 | 0 | 0 | 0 | 0 |
| D-04-3-04 | DAT | D-04 | - | 3 | 4 | yes | Str | - | 2.90 | - | 1 | 0 | 0 | 0 | 0 |
| D-04-3-05 | DAT | D-04 | - | 3 | 5 | yes | Str | - | 3.40 | - | 1 | 0 | 0 | 0 | 0 |
| D-04-3-06 | DAT | D-04 | - | 3 | 6 | yes | Str | - | 3.90 | - | 1 | 0 | 0 | 0 | 0 |
| D-05-3-02 | DAT | D-05 | - | 3 | 2 | yes | Str | - | 1.90 | - | 1 | 0 | 0 | 0 | 0 |
| D-05-3-03 | DAT | D-05 | - | 3 | 3 | yes | Str | - | 2.40 | - | 1 | 0 | 0 | 0 | 0 |
| D-05-3-04 | DAT | D-05 | - | 3 | 4 | yes | Str | - | 2.90 | - | 1 | 0 | 0 | 0 | 0 |
| D-05-3-05 | DAT | D-05 | - | 3 | 5 | yes | Str | - | 3.40 | - | 1 | 0 | 0 | 0 | 0 |
| D-06-2-04 | DAT | D-06 | - | 2 | 4 | no | SNc | - | 4.00 | - | 0 | 0 | 1 | 0 | 0 |
| D-06-2-05 | DAT | D-06 | - | 2 | 5 | no | SNc | - | 4.20 | - | 0 | 0 | 1 | 0 | 0 |
| D-06-3-05 | DAT | D-06 | - | 3 | 5 | yes | SNc | - | 4.20 | - | 0 | 0 | 1 | 0 | 0 |
| D-06-3-07 | DAT | D-06 | - | 3 | 7 | yes | SNc | - | 4.60 | - | 0 | 0 | 1 | 0 | 0 |
| D-07-1-04 | DAT | D-07 | - | 1 | 4 | no | SNc | - | 4.00 | - | 0 | 0 | 1 | 0 | 0 |
| D-08-1-04 | DAT | D-08 | - | 1 | 4 | no | SNc | - | 4.40 | - | 0 | 0 | 1 | 0 | 0 |
| D-08-1-05 | DAT | D-08 | - | 1 | 5 | no | SNc | - | 4.60 | - | 0 | 0 | 1 | 0 | 0 |
| D-08-2-05 | DAT | D-08 | - | 2 | 5 | no | SNc | - | 4.20 | - | 0 | 0 | 1 | 0 | 0 |
| D-08-2-06 | DAT | D-08 | - | 2 | 6 | no | SNc | - | 4.40 | - | 0 | 0 | 1 | 0 | 0 |
| D-08-2-07 | DAT | D-08 | - | 2 | 7 | no | SNc | - | 4.60 | - | 0 | 0 | 1 | 0 | 0 |
| D-08-3-05 | DAT | D-08 | - | 3 | 5 | yes | SNc | - | 4.60 | - | 0 | 0 | 1 | 0 | 0 |
| D-09-1-01 | DAT | D-09 | m | 1 | 1 | no | Str | Str | 2.20 | 2.20 | 1 | 1 | 0 | 0 | 0 |
| D-09-3-01 | DAT | D-09 | m | 3 | 1 | no | Str | Str | 2.90 | 2.90 | 1 | 1 | 0 | 0 | 0 |
| D-09-5-02 | DAT | D-09 | m | 5 | 2 | yes | Str | Str | 2.50 | 2.50 | 1 | 1 | 0 | 0 | 0 |
| D-09-6-01 | DAT | D-09 | m | 6 | 1 | yes | Str | Str | 2.50 | 2.50 | 1 | 0 | 0 | 0 | 0 |
| D-10-1-02 | DAT | D-10 | m | 1 | 2 | no | Str | Str | 2.20 | 2.20 | 1 | 1 | 0 | 0 | 0 |
| D-10-2-01 | DAT | D-10 | m | 2 | 1 | no | Str | Str | 2.50 | 2.50 | 1 | 1 | 0 | 0 | 0 |
| D-11-1-01 | DAT | D-11 | f | 1 | 1 | no | Str | Str | 2.30 | 2.30 | 1 | 1 | 0 | 0 | 0 |
| D-11-1-02 | DAT | D-11 | f | 1 | 2 | no | Str | Str | 2.50 | 2.50 | 1 | 1 | 0 | 0 | 0 |
| D-11-2-01 | DAT | D-11 | f | 2 | 1 | no | Str | Str | 2.70 | 2.70 | 1 | 1 | 0 | 0 | 0 |
| D-11-3-01 | DAT | D-11 | f | 3 | 1 | no | Str | Str | 2.90 | 2.90 | 1 | 1 | 0 | 0 | 0 |
| D-12-3-01 | DAT | D-12 | f | 3 | 1 | no | Str | Str | 2.60 | 2.60 | 1 | 1 | 0 | 0 | 0 |
| D-13-1-02 | DAT | D-13 | f | 1 | 2 | no | Str | Str | 2.60 | 2.60 | 1 | 1 | 0 | 0 | 0 |
| D-13-2-01 | DAT | D-13 | f | 2 | 1 | no | Str | Str | 3.20 | 3.20 | 1 | 1 | 0 | 0 | 0 |
| D-13-3-01 | DAT | D-13 | f | 3 | 1 | no | Str | Str | 2.90 | 2.90 | 1 | 1 | 0 | 0 | 0 |
| D-14-1-01 | DAT | D-14 | f | 1 | 1 | no | Str | Str | 2.50 | 2.50 | 1 | 1 | 0 | 0 | 0 |
| D-14-2-02 | DAT | D-14 | f | 2 | 2 | no | Str | Str | 2.90 | 2.90 | 1 | 1 | 0 | 0 | 0 |
| D-15-4-01 | DAT | D-15 | m | 4 | 1 | no | Str | Str | 3.00 | 3.00 | 1 | 1 | 0 | 0 | 0 |
| D-15-4-02 | DAT | D-15 | m | 4 | 2 | no | Str | Str | 3.30 | 3.30 | 1 | 1 | 0 | 0 | 0 |
| D-15-5-01 | DAT | D-15 | m | 5 | 1 | yes | Str | Str | 2.70 | 2.70 | 0 | 1 | 0 | 0 | 0 |
| D-16-1-01 | DAT | D-16 | m | 1 | 1 | yes | SNc | Str | 3.40 | 1.50 | 0 | 1 | 1 | 0 | 1 |
| D-16-1-02 | DAT | D-16 | m | 1 | 2 | yes | SNc | Str | 3.65 | 1.75 | 0 | 1 | 1 | 0 | 1 |
|  |  |  |  |  |  |  | F1 | F2 | F1 | F2 | F1 | F2 | F1 | F2 |  |
| D-16-3-02 | DAT | D-16 | m | 3 | 2 | yes | SNc | Str | 3.60 | 2.50 | 0 | 0 | 1 | 0 | 0 |
| D-16-4-02 | DAT | D-16 | m | 4 | 2 | yes | SNc | Str | 3.80 | 2.30 | 0 | 0 | 0 | 0 | 1 |
| D-17-1-01 | DAT | D-17 | f | 1 | 1 | yes | SNc | Str | 3.60 | 1.70 | 0 | 1 | 1 | 0 | 1 |
| D-17-1-02 | DAT | D-17 | f | 1 | 2 | yes | SNc | Str | 3.80 | 1.90 | 0 | 1 | 1 | 0 | 1 |
| D-17-1-03 | DAT | D-17 | f | 1 | 3 | yes | SNc | Str | 4.05 | 2.15 | 0 | 1 | 1 | 0 | 1 |
| D-17-2-02 | DAT | D-17 | f | 2 | 2 | yes | SNc | Str | 3.60 | 2.10 | 0 | 1 | 1 | 0 | 1 |
| D-17-2-03 | DAT | D-17 | f | 2 | 3 | yes | SNc | Str | 3.80 | 2.30 | 0 | 1 | 0 | 0 | 1 |
| D-17-2-04 | DAT | D-17 | f | 2 | 4 | yes | SNc | Str | 4.00 | 2.50 | 0 | 1 | 1 | 0 | 1 |
| D-17-2-05 | DAT | D-17 | f | 2 | 5 | yes | SNc | Str | 4.20 | 2.70 | 0 | 1 | 1 | 0 | 1 |
| D-17-3-01 | DAT | D-17 | f | 3 | 1 | yes | SNc | Str | 3.50 | 1.80 | 0 | 1 | 0 | 0 | 1 |
| D-17-3-02 | DAT | D-17 | f | 3 | 2 | yes | SNc | Str | 3.75 | 2.05 | 0 | 1 | 1 | 0 | 1 |
| D-17-3-03 | DAT | D-17 | f | 3 | 3 | yes | SNc | Str | 4.00 | 2.30 | 0 | 1 | 1 | 0 | 1 |
| D-17-3-04 | DAT | D-17 | f | 3 | 4 | yes | SNc | Str | 4.25 | 2.55 | 0 | 1 | 1 | 0 | 1 |
| D-17-4-01 | DAT | D-17 | f | 4 | 1 | yes | SNc | Str | 3.60 | 1.70 | 0 | 0 | 0 | 0 | 1 |
| D-17-4-02 | DAT | D-17 | f | 4 | 2 | yes | SNc | Str | 3.85 | 1.95 | 0 | 1 | 1 | 0 | 1 |
| D-17-4-03 | DAT | D-17 | f | 4 | 3 | yes | SNc | Str | 4.10 | 2.20 | 0 | 1 | 1 | 0 | 1 |

## References

1. Schultz, W., Dayan, P. & Montague, P. R. A neural substrate of prediction and reward. Science (1979) 275, 1593–1599 (1997).

2. Schultz, W., Ruffieux, A. & Aebischer, P. The activity of pars compacta neurons of the monkey substantia nigra in relation to motor activation. Experimental Brain Research 1983 51:3 51, 377–387 (1983).

3. Howe, M. W. & Dombeck, D. A. Rapid signalling in distinct dopaminergic axons during locomotion and reward. Nature 535, 505–510 (2016).

4. da Silva, J. A. et al. Dopamine neuron activity before action initiation gates and invigorates future movements. Nature 554, 244–248 (2018).

5. Fan, D., Rossi, M. A. & Yin, H. H. Mechanisms of action selection and timing in substantia nigra neurons. Journal of Neuroscience 32, 5534–5548 (2012).

6. Coddington, L. T. & Dudman, J. T. The timing of action determines reward prediction signals in identified midbrain dopamine neurons. Nature Neuroscience 2018 21:11 21, 1563–1573 (2018).

7. Engelhard, B. et al. Specialized coding of sensory, motor and cognitive variables in VTA dopamine neurons. Nature vol. 570 509–513 (Nature Publishing Group, 2019).

8. Howe, M. W., Tierney, P. L., Sandberg, S. G., Phillips, P. E. M. & Graybiel, A. M. Prolonged dopamine signalling in striatum signals proximity and value of distant rewards. Nature 500, (2013).

9. Cataldi, S., Lacefield, C., Kumar, G. & Sulzer, D. A dopamine-dependent decrease in dorsomedial striatum direct pathway neuronal activity is required for learned motor coordination. doi:10.1101/2021.06.07.447452.

10. Tsai, H. C. et al. Phasic firing in dopaminergic neurons is sufficient for behavioral conditioning. Science (1979) 324, 1080–1084 (2009).

11. Witten, I. B. et al. Recombinase-driver rat lines: Tools, techniques, and optogenetic application to dopamine-mediated reinforcement. Neuron 72, 721–733 (2011).

12. Saunders, B. T., Richard, J. M., Margolis, E. B. & Janak, P. H. Dopamine neurons create Pavlovian conditioned stimuli with circuit-defined motivational properties. Nature Neuroscience 21, 1072–1083 (2018).

13. Ding, J. B., Guzman, J. N., Peterson, J. D., Goldberg, J. A. & Surmeier, D. J. Thalamic gating of corticostriatal signaling by cholinergic interneurons. Neuron 67, 294–307 (2010).

14. Threlfell, S. et al. Striatal Dopamine Release Is Triggered by Synchronized Activity in Cholinergic Interneurons. Neuron 75, 58–64 (2012).

15. Cachope, R. et al. Selective Activation of Cholinergic Interneurons Enhances Accumbal Phasic Dopamine Release: Setting the Tone for Reward Processing. Cell Reports 2, 33–41 (2012).

16. Liu, C. et al. An action potential initiation mechanism in distal axons for the control of dopamine release. Science (1979) 375, 1378–1385 (2022).

17. Mohebi, A. et al. Dissociable dopamine dynamics for learning and motivation. Nature 570, 65–70 (2019).

18. de Jong, J. W., Fraser, K. M. & Lammel, S. Mesoaccumbal Dopamine Heterogeneity: What Do Dopamine Firing and Release Have to Do with It? (2022) doi:10.1146/annurev-neuro-110920.

19. Lee, S. J. et al. Cell-type-specific asynchronous modulation of PKA by dopamine in learning. Nature | 590, 451 (2021).

20. Patriarchi, T. et al. An expanded palette of dopamine sensors for multiplex imaging in vivo. Nature Methods 2020 17:11 17, 1147–1155 (2020).

21. Petreanu, L. et al. Activity in motor–sensory projections reveals distributed coding in somatosensation. Nature 2012 489:7415 489, 299–303 (2012).

22. Cox, C. L., Denk, W., Tank, D. W. & Svoboda, K. Action potentials reliably invade axonal arbors of rat neocortical neurons. Proc Natl Acad Sci U S A 97, 9724–9728 (2000).

23. Turner, T. J. Nicotine Enhancement of Dopamine Release by a Calcium-Dependent Increase in the Size of the Readily Releasable Pool of Synaptic Vesicles. Journal of Neuroscience 24, 11328–11336 (2004).

24. Woodward, J. J., Judson Chandler, L. & Leslie, S. W. Calcium-dependent and -independent release of endogenous dopamine from rat striatal synaptosomes. Brain Research 473, 91–98 (1988).

25. Tritsch, N. X., Ding, J. B. & Sabatini, B. L. Dopaminergic neurons inhibit striatal output through non-canonical release of GABA. Nature 2012 490:7419 490, 262–266 (2012).

26. Poulin, J.-F. F. et al. Mapping projections of molecularly defined dopamine neuron subtypes using intersectional genetic approaches. Nature Neuroscience 21, 1260–1271 (2018).

27. Howe, M. et al. Coordination of rapid cholinergic and dopaminergic signaling in striatum during spontaneous movement. Elife 8, (2019).

28. Gritton, H. J. et al. Unique contributions of parvalbumin and cholinergic interneurons in organizing striatal networks during movement. Nature Neuroscience 22, 586–597 (2019).

29. Tian, L. et al. Imaging neural activity in worms, flies and mice with improved GCaMP calcium indicators. Nature Methods 6, 875–881 (2009).

30. Poulin, J. F. et al. Defining midbrain dopaminergic neuron diversity by single-cell gene expression profiling. Cell Reports 9, 930–943 (2014).

31. Poulin, J. F., Gaertner, Z., Moreno-Ramos, O. A. & Awatramani, R. Classification of Midbrain Dopamine Neurons Using Single-Cell Gene Expression Profiling Approaches. Trends in Neurosciences vol. 43 155–169 (2020).

32. Fenno, L. E. et al. Comprehensive Dual- and Triple-Feature Intersectional Single-Vector Delivery of Diverse Functional Payloads to Cells of Behaving Mammals. Neuron 107, 836–853.e11 (2020).

33. Madisen, L. et al. Transgenic mice for intersectional targeting of neural sensors and effectors with high specificity and performance. Neuron 85, 942–958 (2015).

34. Farassat, N. et al. In vivo functional diversity of midbrain dopamine neurons within identified axonal projections. Elife 8, 1–27 (2019).

35. Lerner, T. N. et al. Intact-Brain Analyses Reveal Distinct Information Carried by SNc Dopamine Subcircuits. Cell 162, 635–647 (2015).

36. Lammel, S., Ion, D. I. I., Roeper, J. & Malenka, R. C. C. Projection-Specific Modulation of Dopamine Neuron Synapses by Aversive and Rewarding Stimuli. Neuron 70, 855–862 (2011).

37. Parker, N. F. et al. Reward and choice encoding in terminals of midbrain dopamine neurons depends on striatal target. (2016) doi:10.1038/nn.4287.

38. Fallon, J. H. & Moore, R. Y. Catecholamine innervation of the basal forebrain IV. Topography of the dopamine projection to the basal forebrain and neostriatum. Journal of Comparative Neurology 180, 545–579 (1978).

39. Mendonça, M. D. et al. Transient dopamine neuron activity precedes and encodes the vigor of contralateral movements. bioRxiv 2021.04.20.440527 (2021) doi:10.1101/2021.04.20.440527.

40. Pereira Luppi, M. et al. Sox6 expression distinguishes dorsally and ventrally biased dopamine neurons in the substantia nigra with distinctive properties and embryonic origins. Cell Reports 37, 109975 (2021).

